# Macrophage signaling and function are regulated by distinct sterol biochemistries

**DOI:** 10.1101/2025.08.07.669025

**Authors:** Jazmine D. W. Yaeger, Jason G. Kerkvliet, Bijaya Pradhan, Amelia G. Lawver, Sonali Sengupta, Natalie W. Thiex, Kevin R. Francis

## Abstract

Membranes require continuous reorganization of lipid components, including sterols, to dynamically alter their rigidity to deform and bend during scission events which occur during fundamental cellular functions such as endocytosis. While diseases of cholesterol biosynthesis result in reduced cellular cholesterol and accumulation of precursor sterols, limited studies have addressed the intracellular consequences of disease-associated sterol changes on the ability of eukaryotic cellular membranes to function and signal normally. Here, we utilized bone marrow-derived macrophages (BMDMs) to investigate how altered sterol content impacts macrophage signaling and membrane function. Through pharmacological inhibition of cholesterol biosynthetic enzymes, reduced cholesterol and increased levels of disease-associated sterol intermediates coincided with reduced expression of cell surface proteins and impaired macropinocytosis. Macropinocytic activity was sensitive to both reduced plasma membrane cholesterol and sterols containing functional groups substituted for the C3 hydroxyl group. Transcriptomic analyses of cholesterol-inhibited BMDMs revealed alterations in immune and chemokine signaling pathways. Decreased cholesterol was also associated with dysregulated vesicular sorting pathways and elevated expression of endosomal/lysosomal markers. Disrupted endosome expression and impaired macropinocytosis was also observed in BMDMs from mouse models of the cholesterol biosynthesis disorder Smith-Lemli-Opitz syndrome (SLOS). Our findings detail an important connection between sterol imbalance, membrane dynamics, and immune cell function.

## INTRODUCTION

Sterols are a fundamental component of cellular structure and function where they support plasma membrane and organelle membrane function, secure proteins and carbohydrates to specific subcellular loci, serve as a foundation for steroid production, and activate signaling pathways to regulate metabolic processes. Cholesterol, the most abundant sterol in mammalian systems, maintains an amphipathic structure which may undergo modifications such as esterification to enhance its stability in hydrophobic cellular compartments. Sterol intermediates are often short-lived molecules in mammalian cells, as they are rapidly converted through enzymatic activity toward cholesterol. While exceptions exist, like the accumulation of desmosterol in testes (1) or the developing brain (2), accumulation of sterol intermediates is typically associated with disease.

Rare genetic disorders of post-squalene cholesterol biosynthesis result in an often-dramatic reduction in cholesterol levels and aberrant accumulation of sterol intermediates (3). The most prevalent of these diseases is Smith-Lemli-Opitz syndrome (SLOS), stemming from a mutation in the *DHCR7* gene and leading to elevated levels of 7-dehydrocholesterol (7DHC) in-parallel with decreased cholesterol (4, 5). Other cholesterol biosynthesis disorders result in the accumulation of 4,4-dimethyl sterols (6), zymostenol/zymosterol (7), lathosterol (8), or desmosterol (9). Clinical symptoms within these disorders are diverse (10), suggesting the intermediate sterols themselves likely contribute to pathology. However, the impacts of cholesterol versus sterol intermediates at the cellular level has not been fully explored.

Cellular membranes require complex associations between various lipid species and proteins to function normally. Cholesterol shuttles to and from the plasma membrane (11) through membrane bending, vesicle fusion, and vesicular and non-vesicular transport processes (12). Alterations in lipid composition and redistribution of membrane lipids allow for changes in membrane shape and mobility, including within monocyte derivatives (13). Cholesterol homeostasis plays a critical part in defining a cell’s immune-associated identity (14–16). For example, disruption of the cholesterol biosynthetic pathway induces a type I interferon response (17). Also, the reorganization of cholesterol pools is a critical adaptation in regulating macrophage defense mechanisms (18).

Macrophages, prominent immune cells existing in specialized forms in nearly all tissue types, require dramatic membrane changes for cellular functions, including engulfment and breakdown of extracellular products. As resident immune cells, macrophages assess changes in environmental stimuli through pattern recognition receptors (19) and changes in transcriptional programming leading to a spectrum of cellular responses ranging from pro-inflammatory to anti-inflammatory. In addition to cholesterol, macrophages also utilize additional sterols. Lanosterol levels increase in macrophages during activation of toll-like receptor 4 (20). Cholestenone accumulation coincides with infection of *Myobacterium tuberculosis* (21). Microglia isolated from *Dhcr7*^T93M/Δ3-5^ mice which model SLOS exhibit pro-inflammatory phenotypes (22). While these studies highlight an essential link between sterol metabolism and macrophage immune function, questions remain regarding sterol-specific impacts on immune cell response and function.

In this study, we utilized pharmacological and genetic models of cholesterol biosynthesis disorders to discern the effects of cholesterol reduction versus sterol intermediate accumulation on bone marrow-derived macrophage (BMDM) signaling and function. We show that disruption of BMDM cholesterol biosynthesis not only alters macrophage biochemistry, but also disrupts proliferation, surface marker expression, endocytic functions, chemotactic signaling, and vesicular trafficking pathways. We further demonstrate a link between immune activation and cholesterol metabolism in macrophages. These results further delineate the significant role of sterol biochemistry within macrophages and detail disease-relevant, sterol-specific impacts on macrophage cellular processes and function.

## MATERIALS AND METHODS

### Animals and housing

Mouse lines used for these studies include C57BL/6J (The Jackson Laboratory, Bar Harber, ME), *Dhcr7*^T93M/+^, and *Dhcr7*^T93M/Δ3-5^. By breeding mice of *Dhcr7*^Δ3-5/+^ and *Dhcr7*^T93M/T93M^ backgrounds (kindly provided by Dr. Forbes Porter, Eunice Kennedy Shriver National Institute of Child Health and Human Development), *Dhcr7*^T93M/Δ3-5^ mice were generated. Heterozygous *Dhcr7*^T93M/Δ3-5^ mice carry a deletion of exons III, IV, and a portion of V on one allele, as well as a dinucleotide mutation in codon 89 of the *Dhcr7* gene on the other allele (23, 24). Breeding pairs for each genetic line were formed from mice at least 2 months of age and breeders were retired by 9 months of age. All animals were kept on a C57BL/6 background and housed in the animal resource center at Sanford Research or South Dakota State University on a 12 h/12 h light-dark cycle. Mice were fed a standard diet *ad libitum*. Animal resource centers are access-controlled environments and managed by trained technicians and veterinarians who perform daily health checks. Macrophages utilized for experiments were isolated and cultured from male 6-week-old mice. Animal work was evaluated and approved by the Institutional Animal Care and Use Committees at Sanford Research (protocol # 2023-0091) and South Dakota State University (protocol # 2402-027A).

### Bone marrow-derived macrophage isolation and cell culture

Bone marrow-derived macrophages (BMDM) were isolated from 6-week-old male mice (supplemental Fig. S1A) using previously described methods (25). Mice were euthanized by CO_2_ asphyxiation followed by cervical dislocation. Sacrificed animals were sterilized with 70% ethanol and incisions were used to expose the hind limbs. Muscle was removed and femurs were carefully dissected, washed in sterile phosphate buffer saline (PBS), and kept on ice. The apical ends of the isolated femurs were removed and the bone marrow was flushed using PBS into a sterile conical tube. Collected bone marrow was centrifuged for 5 min at 300 x g, the pellet was suspended in media and plated onto non-tissue culture treated dishes. After 48 h, without removing any of the previously supplied culture media, fresh media was added. Mature BMDMs were found adhered to the dish after another 48 h. Culture medium for BMDMs includes DMEM (ATCC, Cat. 30-2002), 20% (v/v) fetal bovine serum (FBS; Corning), 0.1% (v/v) penicillin/streptomycin (Life Technologies, 15140122), 50 ng/mL colony-stimulating factor 1 (CSF1; BioLegend, 574804), and 57.6 nM β-mercaptoethanol (Life Technologies, 21985023). Media was equilibrated to pH 7.4 by placing it in an incubator set to 37°C and 5% CO_2_ prior to use. To induce *de novo* cholesterol biosynthesis in macrophages, cells were washed gently with PBS and cultured in 20% (v/v) lipoprotein-deficient serum (LPDS) for 48 h.

### Preparation of lipoprotein-deficient serum (LPDS)

Fetal bovine serum (FBS) in absence of neutral lipids including sterols and triglycerides (called lipoprotein-deficient serum, LPDS) was produced from techniques described previously (26–29). Briefly, oxidation initiated by trace peroxides was avoided by adding 0.1 mg ethylenediamine tetraacetate (EDTA) for every 50 mL of FBS. An organic phase mixture (3:2 ratio diisopropyl ether:n-butanol) was combined with the serum (1:2 ratio) and stirred protected from light for 1 h. After stirring, the organic phase was discarded while the remaining solution was centrifuged at 4°C for 15 min at 2200 rpm. The aqueous layer was filtered and freeze-dried to remove residual organic solvents. The lyophilized LPDS product was resuspended in molecular grade H_2_O and supplemented with insulin-transferrin-selenium (ITS-G; Gibco, 41400045). The final LPDS product was filter-sterilized, aliquoted, and stored at -20°C. Sterol content in LPDS batches was assessed using gas chromatography paired with mass spectrometry (GC-MS).

### Gas chromatography-mass spectrometry (GC-MS)

GC-MS was performed as previously described (30, 31). BMDM cell pellets were flash frozen on dry ice and stored at -20°C. Pellets were resuspended in 1 mL H_2_O and lysed by successive freeze-thaw cycles. 50 µL of cell lysate was used for protein quantification (Micro BCA Protein Assay Kit, Thermo Fisher Scientific, 23235), while the remainder was combined with 1 mL saponification buffer (92% ethanol, 7% KOH, and 10 µg/mL coprostan-3-ol as an internal standard) and heated at 60°C for 1 h. After saponification, 1 mL of H_2_O was added to each sample and aqueous phase separation was initiated by mixing in 3 mL ethyl acetate. The samples were centrifuged for 5 min at 2200 rpm and the organic phase was mixed with 2 mL of H_2_O. After a second centrifugation, the top layer of lipids was isolated and dehydrated at 50°C under a constant flow of nitrogen gas. Dried sterols were dissolved in 50 µL pyridine and derivatized with 50 µL N,O-bis(trimethylsilyl)trifluoroacetamide with 1% trimethylchlorosilane (BSTFA + 1% TMCS, Thermo Fisher Scientific, TS-38831) for 1 h at 50°C. 1 µL of derivatized sterol samples were injected into a split injection port (4 mm ID x 78.5 mm quartz wool liner, Restek 23309) on an Agilent 7890 gas chromatograph housed with a 0.18 mm ID x 20 m 1,4-bis(dimethylsiloxy)phenylene dimethyl polysiloxane column (Restek, 43602). Helium was used as the carrier gas and was set to a flow rate of 46.9 cm/sec. The GC method utilized proceeded as follows: 170°C for 30 sec, oven temperature was raised to 250°C at a rate of 18°C/min, then increased to 280°C at a rate of 3°C/min, before being held for 7 min at 320°C once the temperature was reached at a rate of 20°C/min. An Agilent 5977B mass spectrometer was set to electron impact mode (70 eV) where it was kept at a source temperature of 275°C. Derivatized ethers of sterols were identified by comparing chromatogram elution times and MS spectra to standards for cholesterol, lathosterol, 7-dehydrocholesterol (7DHC), and desmosterol (Avanti Polar Lipids Inc.). MS spectra were further compared to those accessible through a National Institute of Standards and Technologies Standard Reference Database. Representative spectra fragmentation patterns for samples are available upon request. Sterol abundance was normalized to both an internal standard (coprostan-3-ol) and protein concentration (Micro BCA Protein Assay Kit, Thermo Fisher Scientific, 23235). Data was analyzed using MassHunter software and normalized data was graphed relative to control samples using GraphPad Prism software.

### Small molecule targeting of cholesterol and lipid metabolism

Cholesterol biosynthesis in BMDMs was antagonized through administration of enzyme-targeted small molecule inhibitors (**Fig. 1A**). Macrophages were washed with PBS and incubated with pharmacological agents dissolved in dimethyl sulfoxide (DMSO) and brought to desired concentrations in LPDS media. Cells were treated for 48 h with small molecule inhibitors or vehicle (DMSO) prior to downstream assays. Pharmacological treatments used included simvastatin (inhibitor of 3-hydroxy-3-methyl-glutaryl-coenzyme A reductase, HMG-CoA reductase; 1 µM; Cayman Chemical, 10010344), 17α-OH-progesterone (17α-OHP, inhibitor of sterol-C4-methyl oxidase, SC4MOL; 25 µM; Cayman Chemical, 33154), TASIN-1 (inhibitor of emopamil binding protein, EBP; 1 µM; Cayman Chemical, 2155), AY9944 (inhibitor of 7-dehydrocholesterol reductase, DHCR7; 1 µM; Cayman Chemical, 14611), and U18666A (inhibitor of 24-dehydrocholesterol reductase, DHCR24; 20 nM; Cayman Chemical, 10009085). Indicated dosages used were selected following dose-response analyses of BMDM biochemical sensitivity to each compound with GC-MS. For select assays, the acyl-CoA:cholesterol acyltransferase inhibitor Avasimibe (5 µM) was used to target lipid storage mechanisms. In specified experiments, an inflammatory response was induced by co-treating macrophages from C57BL/6 mice for 48 h with interferon-γ (IFNγ; 2.5 ng/mL) and lipopolysaccharide (LPS; 0.5 µg/mL), followed by co-treatment of IFNγ/LPS with AY9944 for another 48 h. For lipid overload experiments, macrophages were supplemented with oleic acid (OA; Sigma Aldrich, O1383) carried in bovine serum albumin (BSA; Sigma Aldrich, A8806). For lipid challenge experiments, 200 µM OA or vehicle (0.2% BSA) were added to cells for 24 h.

**Figure 1.**
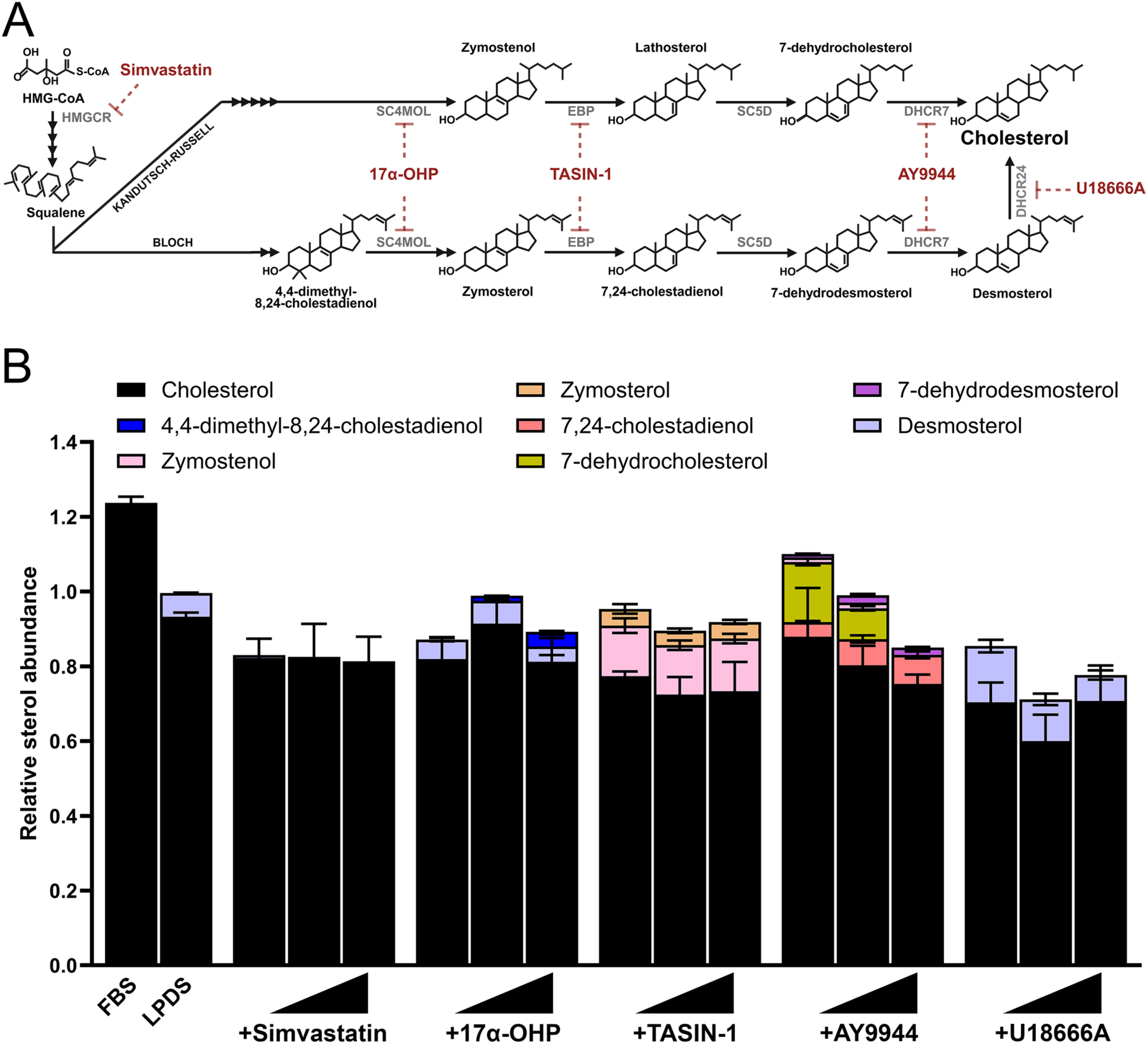
Inhibition of sterol synthesis induces biochemical changes in macrophages. (A) Cholesterol biosynthesis is composed of multiple reactions catalyzed by enzymes (gray text) which can be targeted with small molecule inhibitors (red text). (B) Quantified GC-MS analyses of BMDMs treated for 48 h with small molecule inhibitors of cholesterol biosynthesis show reduced cholesterol and accumulation of sterol precursor molecules (mean ± SEM; n = 3 biological replicates from 3 independent experiments).

### Live-dead assay

To assess cell death caused by impaired cholesterol biosynthesis, we utilized a live-dead kit (Biotium Live-or-Dye Fixable Viability Staining Kit, Cat. 32005) according to manufacturer’s instructions. BMDMs were grown on 12 mm fibronectin-coated (0.25%; Sigma Aldrich, Cat. FC010) coverslips. After 48 h treatment with small molecule inhibitors, cells were washed with PBS and incubated cells for 30 min protected from light in culture medium containing a 1:1000 dilution of Fixable Dead Cell Dye. As a positive control for cell death, untreated cells were incubated in a 15% ethanol/85% PBS solution for 10 min. Cells were rinsed and fixed for 15 min with 4% paraformaldehyde (PFA; Electron Microscopy Sciences, 15714). After rinsing, cells were then permeabilized with 0.05% Triton X-100 (Sigma-Aldrich, 93443) for 10 min and incubated overnight at 4°C with a mouse anti-Ki67 antibody (1:400; Cell Signaling Technologies, Cat. 9449) in blocking buffer (0.05% TritonX-100; 5% goat serum, Jackson ImmunoResearch Laboratories, Cat. 005-000-121). Following rinsing, cells were incubated for 1 h with anti-rabbit AlexaFluor-Cy5 (1:500; Life Technologies, A11001) and Hoechst 33342 nuclear marker (1:10,000; Invitrogen, H3570). Images were captured using a Nikon CSU-W1 Spinning Disk Super Resolution by Optical Pixel Reassignment (SoRa) system, equipped with NIS-Elements analysis software (Nikon) and analyzed for percent positive cells for the live-dead stain and Ki67 using QuPath image analysis software (version 0.2.3).

### Immunocytochemistry and image acquisition

BMDMs were seeded and grown on 12 mm fibronectin-coated (0.25%) glass coverslips for 24 h prior to drug treatment (described above). Cells were fixed in 4% PFA for 15 min, washed with PBS, and permeabilized in 0.05% Triton X-100 for 15 min. Blocking buffer consisting of 0.05% Triton X-100 and 5% goat serum in PBS was added for 1 h at room temperature to coverslips. Permeabilized cells were incubated with primary antibodies in blocking buffer at 4°C overnight. The primary antibodies used include rabbit anti-Rab7 (Cell Signaling Technologies, 9367, 1:200), rat anti-CD68 (Bio-Rad, MCA1957, 1:500), and rabbit anti-perilipin-2 (Novus, NB110-40877, 1:200). After primary antibody incubation, cells were washed three times with PBS before being exposed to species-appropriate Alexa Fluor-conjugated secondary antibodies (Life Technologies, A11001, A21437 and A31570; 1:500) for 1 h incubation at room temperature protected from light. Where indicated, cells were stained with phalloidin conjugated to a far-red Alexa 647 dye (Thermo Fisher Scientific, A22287, 1:400), BODIPY 505/515 neutral lipid stain (Invitrogen, H3570, 1:2000), and Hoechst 33342 nuclear counterstain (Invitrogen, H3570, 1:10,000). Image acquisition was performed using Nikon CSU-W1 Spinning Disk Super Resolution by Optical Pixel Reassignment (SoRa) system, equipped with NIS-Elements analysis software (Nikon). Image analysis was completed using Fiji software (ImageJ 1.54f).

### Flow cytometry and endocytosis assays

Accuri C6 Plus (BD Biosciences) or FACS Jazz (BD Biosciences) flow cytometers were used. Cytometers were cleaned (BD FACS Cleaning Solution, BD Biosciences, 34035) and rinsed with ddH_2_O prior to use. Lasers were aligned with tracking beads (BD Biosciences, 661414) to ensure system consistency. For surface marker expression, BMDMs were harvested from non-tissue culture treated dishes by incubating at 4°C for 10 min in cold PBS. Cells were pelleted by centrifugation at 300 x g for 5 min and suspended in surface marker antibodies diluted in cold 1% FBS in PBS. Cells were incubated with antibodies on ice for 15 min. Cells were then washed three times with cold 1% FBS in PBS solution before being suspended in 250 µL for analysis. Antibodies used for surface marker analysis included CD11b (Fisher Scientific, BDB557672), CD11c (BioLegend, 117313), CD16/32 (BioLegend, 101325), CD80 (BioLegend, 104707), and CD115 (Fisher Scientific, 50-112-8909). Isotype controls (BioLegend, 400612, 400608, 600625) were analyzed in-parallel to account for non-specific binding. Endocytosis assays used lucifer yellow CH, lithium salt (LY; Fisher Scientific, L453) to measure macropinocytosis and Texas Red dextran, 70,000 MW (Fisher Scientific, D1864) uptake for assessment of clathrin-mediated endocytosis. Following treatment, BMDMs were incubated in DMEM containing 200 ng/mL CSF1 (for macrophage stimulation) and 0.5 mg/mL LY or 0.25 mg/mL Texas red dextran for 1 h at 37°C and 5% CO2. Cells were then rinsed three times with PBS, collected, and analyzed. For each biological replicate, BMDMs were kept on ice during LY or Texas red dextran exposure as a negative control to inhibit endocytosis. Viability stains (NucRed, Invitrogen, R37113; 7-AAD, BioLegend, 420403) were used to gate for live cell populations.  ≥ 10,000 events were recorded per sample and analyzed with FlowJo and GraphPad Prism.

### Cholesterol stripping and sterol loading

Sterol stripping and loading experiments were carried out as previously described (27) with slight modifications. Methyl-β-cyclodextrin (MβCD; 5 mM, Sigma, C4951) was suspended in serum-free DMEM. Sterols used include cholesterol (Avanti Polar Lipids, 700000P), 7DHC (Avanti Polar Lipids, 70066P), desmosterol (Avanti Polar Lipids, 700060P), cholestenone (Avanti Polar Lipids, 700065P), and cholesterol sulfate (Avanti Polar Lipids, 700016). Sterols were resuspended in a 1:1 chloroform:methanol mix to 50 mg/mL. The desired amount of sterol (1:7 molar ratio) was concentrated to dryness under constant flow of nitrogen and diluted in 5 mM MβCD-DMEM solution. MβCD-sterol solutions were sonicated for 5 min, incubated overnight at 37°C with continuous agitation, 0.45 µm filtered, and stored at 4°C. For direct loading of sterols into BMDMs, cells were incubated with 5 mM empty MβCD for 1 h at 37°C/5% CO_2_) then MβCD-sterol solutions containing 0.5 mg/mL LY and 200 ng/mL CSF1 for another 1 h. Cells were washed three times, collected and analyzed by flow cytometry.

### Whole genome mRNA sequencing

BMDMs isolated from 6-week, male C57BL/6 mice and treated for 48 h in FBS, LPDS, simvastatin, AY99944, and U18666A conditions before cell pellets were harvested and stored in RNA later. RNA extraction was performed by the Functional Genomics and Bioinformatics core at Sanford Research. Cell pellets were homogenized with a 21G needle in Buffer RLT (Qiagen, 79216) and RNA extraction with on-column DNase digestion performed using the RNeasy mini kit (Qiagen, 74104) following manufacturer’s instructions. RNA was quantified using a NanoDrop spectrophotometer and quality was assessed with a Bioanalyzer using a RNA 6000 Nano Kit (Agilent Technologies, 50671511). cDNA library preparation from mRNA utilizing random hexamer primers, sequencing using the Illumina Novaseq 6000 platform, read mapping to the reference genome using HISAT2, and identification of differentially expressed genes by DESeq2 were performed by Novogene (Sacramento, CA, USA). Transcripts exhibiting log_2_ fold change ≥ ± 0.5 and *p* < 0.05 between two groups were considered differentially expressed. Pathway analyses and identification of upstream regulators were performed using Ingenuity Pathway Analysis software. Gene Ontology (GO) enrichment analysis was performed using clusterProfiler R Package (32) by the Functional Genomics and Bioinformatics core.

### Quantitative real-time polymerase chain reaction (qRT-PCR)

Cell pellets were collected and flash frozen in liquid nitrogen. Total RNA was isolated using the Total RNA Kit (Omega BioTek, R6834-01) and quality validated with a NanoDrop Lite spectrophotometer (Thermo Fisher Scientific). cDNA was synthesized from 1 µg total RNA using the High-Capacity cDNA Reverse Transcription Kit (Applied Biosystems, 4368814) according to manufacturer’s protocols. qRT-PCR was performed using SYBR green chemistry (Luna Universal SYBR green qPCR mix, New England Biolabs, M3003X) on a CFX-96 real-time PCR system (Bio-Rad Laboratories) following manufacturer’s protocols. Fold change was calculated using ddCT method normalized to *Gapdh* and *Actb*. Primers used include (shown 5’ to 3’): *Hmgcr* forward, 5’-AGCTTGCCCGAATTGTATGTG-3’; *Hmgcr* reverse, 5’-TCTGTTGTGAACCATGTGACTTC-3’; *Dhcr7* forward, 5’-AGGCTGGATCTCAAGGACAAT-3’; *Dhcr7* reverse, 5’-GCCAGACTAGCATGGCCTG-3’; *Dhcr24* forward, 5’-CTCTGGGTGCGAGTGAAGG-3’; *Dhcr24* reverse, 5’-TTCCCGGACCTGTTTCTGGAT-3’; *Gapdh* forward, 5’-AGGTCGGTGTGAACGGATTTG-3’; *Gapdh* reverse, 5’-TGTAGACCATGTAGTTGAGGTCA-3’; *Actb* forward, 5’-GGCTGTATTCCCCTCCATCG-3’, *Actb* reverse, 5’-CCAGTTGGTAACAATGCCATGT-3’.

### Accession codes

RNA sequencing data are available in the Gene Expression Omnibus (GEO) database (http://www.ncbi.nlm.nih.gov/gds) under the accession number GSE300253.

### Statistical analysis

GraphPad Prism 8.0.2 (GraphPad Software, Inc., CA, US) was utilized for all statistical analyses. The Brown-Forsythe test was used to test for homogeneity of variances. When variances were equal, data was analyzed using one-way ANOVA and Dunnett’s *post hoc* test relative to vehicle or LPDS control group. When variances were unequal, Welch’s ANOVA and *post hoc* Dunnett’s T3 test were utilized. Significance was accepted as *p* < 0.05. All statistical details for each experiment can be found within the figure legends.

## RESULTS

### Disruption of cholesterol biosynthesis alters the macrophage sterol profile

To analyze sterol impacts on macrophage biology, distinct enzymatic steps within the cholesterol biosynthetic pathway can be inhibited through pharmacological antagonists when applied in lipid-depleted environmental conditions (**Fig. 1A**). Removal of environmental lipids through BMDM culture in lipoprotein-deficient serum (LPDS) reduced cellular cholesterol levels by ∼35% (**Fig. 1B**, supplemental Fig. S1B, S1C), suggesting BMDMs require internalization of environmental cholesterol to maintain homeostatic levels of cholesterol. qRT-PCR analysis demonstrated exposure to 20% LPDS conditions induced transcription of key cholesterol biosynthesis genes (supplemental Fig. S1D-F), indicating BMDMs attempt to adapt to cholesterol-depleted environmental conditions. For example, increased transcription of *Dhcr24* (supplemental Fig. S1F) coincided with accumulation of desmosterol in LPDS environment (**Fig. 1B**, supplemental Fig. S1B). Targeting the cholesterol biosynthesis with simvastatin, an HMG-CoA reductase inhibitor, both reduced cholesterol levels and eliminated LPDS-induced accumulation of desmosterol (**Fig. 1B**), an effect that was simvastatin dose independent (**Fig. 1B**). 17α-hydroxyprogesterone (17α-OHP), an SC4MOL antagonist, induced BMDM accumulation of 4,4-dimethyl-8,24-cholestadienol, but still showed moderate levels of desmosterol (**Fig. 1B**). TASIN-1, an EBP inhibitor, resulted in modest amounts of zymostenol and zymosterol in BMDMs (**Fig. 1B**). The Dhcr7 antagonist AY9944 produced accumulation of 7-dehydrocholesterol (7DHC), 7,24-cholestadienol, zymostenol, and 7-dehydrodesmosterol (**Fig. 1B**). Inhibition of Dhcr24 with U18666A strongly attenuated cholesterol synthesis, reducing cholesterol levels by ∼20% compared to LPDS alone (**Fig. 1B**). Elevated levels of desmosterol were observed as expected after U18666A treatment (**Fig. 1B**). These results demonstrate that biochemical changes arise in BMDMs upon cholesterol-deficient culture and suggest BMDMs may physiologically adapt to attempt to maintain a threshold level of cholesterol.

Since inhibition of cholesterol biosynthesis shifted sterol profiles in macrophages (**Fig. 1**), we next asked if these alterations impacted cell viability or proliferation. Cell viability assays showed inhibition of cholesterol biosynthesis was not toxic to macrophages (supplemental Fig. S2A, S2B). Analyses for Ki67, expressed in actively dividing cells, showed a reduction after 17α-OHP, TASIN-1, or AY9944 treatment; however, treatments which maintained desmosterol levels (simvastatin, U18666A, or avasimibe) did not impact proliferation (supplemental Fig. S2A, S2C, S2D). These data suggest that maintenance of macrophage proliferative capacity in cholesterol-depleted culture, may proceed through a sterol-specific mechanism.

### The sterol profile of macrophages regulates their differentiation, activation and function

Immune response activation was previously found in macrophages upon genetic deletion of SREBP cleavage-activating protein, inhibiting expression of cholesterol and fatty acid biosynthesis genes (17, 18).

To determine if cholesterol reduction or the accumulation of precursor sterols is related to the immunogenic state of BMDMs, we characterized the impact of cholesterol biosynthesis inhibition on expression of immune cell surface markers with flow cytometry (**Fig. 2A, 2B**). Expression of CD11b (also known as integrin alpha M, Mac-1, complement receptor 3), an integrin important for cell-to-cell adhesion and matrix attachment (33, 34), was reduced by LPDS culture alone as well as LPDS culture plus treatment with TASIN-1, AY9944, or U18666A (**Fig. 2C**). Expression of the integrin CD11c (35) was reduced by LPDS culture and further inhibited by TASIN-1 (**Fig. 2D**). Expression of CD16/32, the FcgII and FcgIII phagocytic receptors (36, 37), was inhibited by culture in LPDS and further inhibited by AY9944 (**Fig. 2E**). Although surface expression of the co-stimulatory molecule CD80 (38) was significantly reduced in LPDS, further antagonization of cholesterol biosynthesis with inhibitors had no additive effect (**Fig. 2F**). Expression of the CSF1 receptor CD115 is critical for differentiation of monocytes to macrophages and chemotactic migration of macrophages (39, 40), was specifically inhibited by AY9944 relative to LPDS conditions (**Fig. 2G**). Overall, depletion of lipids from the culture medium and inhibition of cholesterol biosynthesis impaired cell surface expression of critical proteins regulating macrophage differentiation, activation and function. These findings suggest that cholesterol deficient macrophages may have impaired immune function, possibly in a sterol specific manner.

**Figure 2.**
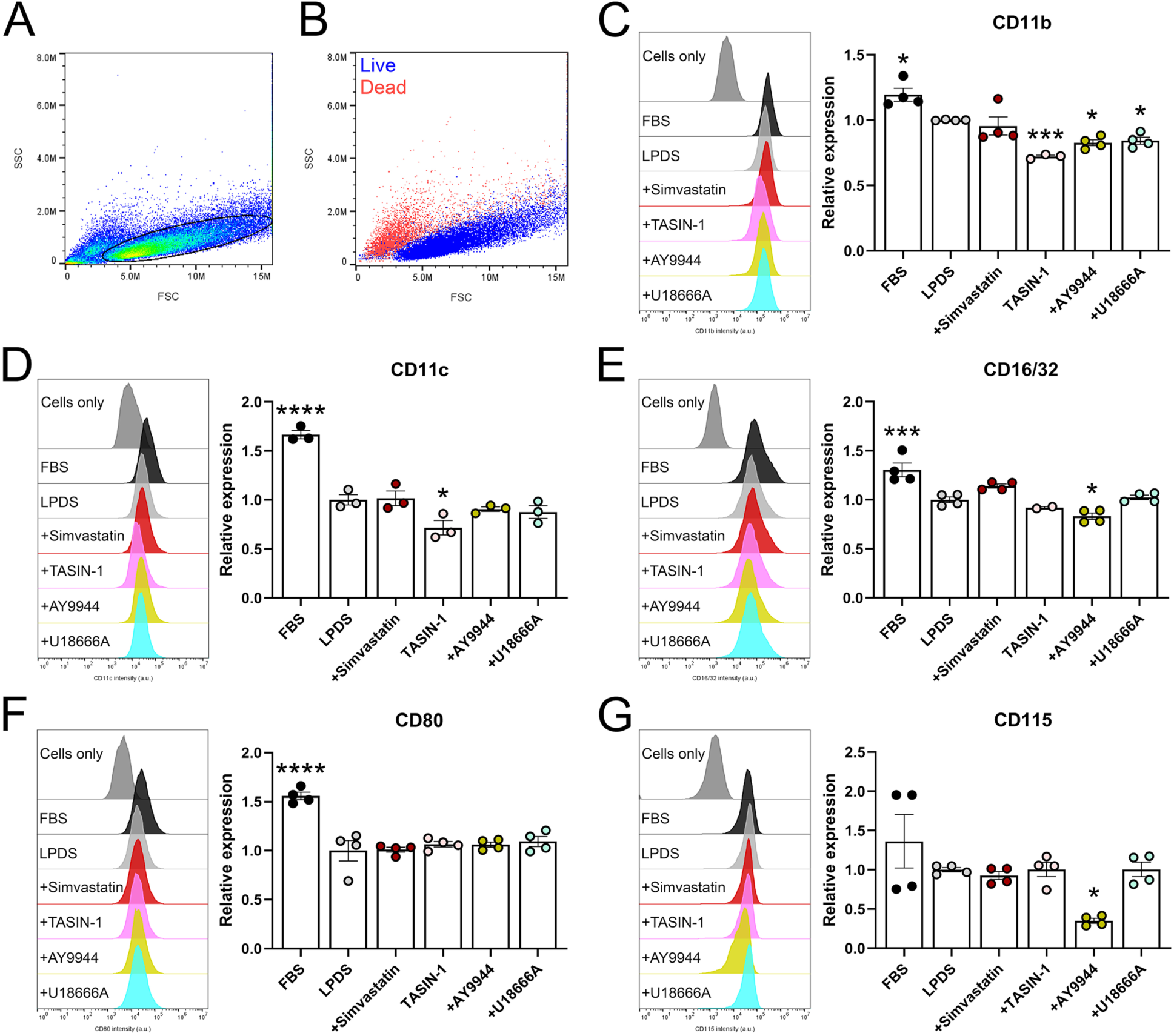
Macrophage surface proteins are altered by sterol change. (A) Representative distribution of forward (FSC) and side scatter (SSC) events showing gating position for detecting live BMDMs with flow cytometry. (B) Representative distribution of live and dead populations of BMDMS used for flow cytometry analyses. (C) Cell surface expression of CD11b in cholesterol biosynthesis inhibited BMDMs (mean ± SEM; n = 3-4 biological replicates from 4 independent experiments). One-way ANOVA (F_5,17_ = 16.66, p < 0.0001) with Dunnett’s multiple comparisons test (*p ≤ 0.05; ***p ≤ 0.001). (D) Surface expression of CD11c n cholesterol biosynthesis inhibited BMDMs (mean ± SEM; n = 3 biological replicates from 3 independent experiments). One-way ANOVA (F_5,12_ = 32.35, p < 0.0001) with Dunnett’s multiple comparisons test (*p ≤ 0.05; ****p < 0.001). (E) CD16/32 surface expression in BMDMs upon cholesterol biosynthesis inhibition (mean ± SEM; n = 2-4 biological replicates from 4 independent experiments). One-way ANOVA (F_4,15_ = 19.65, p < 0.0001) with Dunnett’s multiple comparisons test (*p ≤ 0.05; ***p ≤ 0.001). (F) CD80 surface expression in BMDMs targeted for cholesterol biosynthesis inhibition (mean ± SEM; n = 4 biological replicates from 4 independent experiments). One-way ANOVA (F_5,18_ = 15.82, p < 0.0001) with Dunnett’s multiple comparisons test (****p < 0.001). (G) Surface expression of CD115 in BMDMs in cholesterol replete compared to cholesterol inhibited conditions (mean ± SEM; n = 4 biological replicates from 4 independent experiments). One-way ANOVA (F_5,18_ = 4.706, p ≤ 0.0064) with Dunnett’s multiple comparisons test (*p ≤ 0.05).

### Macrophages exhibit functional defects in endocytic processes upon sterol synthesis inhibition

Due to the observed changes in macrophage sterol profile and surface marker expression after cholesterol biosynthesis inhibition, we predicted sterol-compromised macrophages to have defects in membrane-associated functions (supplemental fig. S3). To test this, we assayed the ability of BMDMs for their ability to perform macropinocytosis by measuring Lucifer yellow (LY) uptake (41) (**Fig. 3A**). Culture in LPDS conditions alone limited LY uptake (**Fig. 3B, 3C**). The addition of cholesterol biosynthesis inhibitors reduced LY internalization even further (**Fig. 3B, 3C**). We previously demonstrated clathrin-mediated endocytosis was inhibited by cholesterol biosynthesis inhibition in a sterol-specific manner (27). We also measured uptake of Texas red dextran, which is primarily internalized via the mannose receptor in BMDM (41). Texas red dextran uptake was reduced in cholesterol-deficient media and upon cholesterol biosynthesis inhibition (supplemental Fig. S3A, S3B). These results highlight the importance of cholesterol homeostasis for the function of the plasma membrane and maintenance of macrophage function.

**Figure 3.**
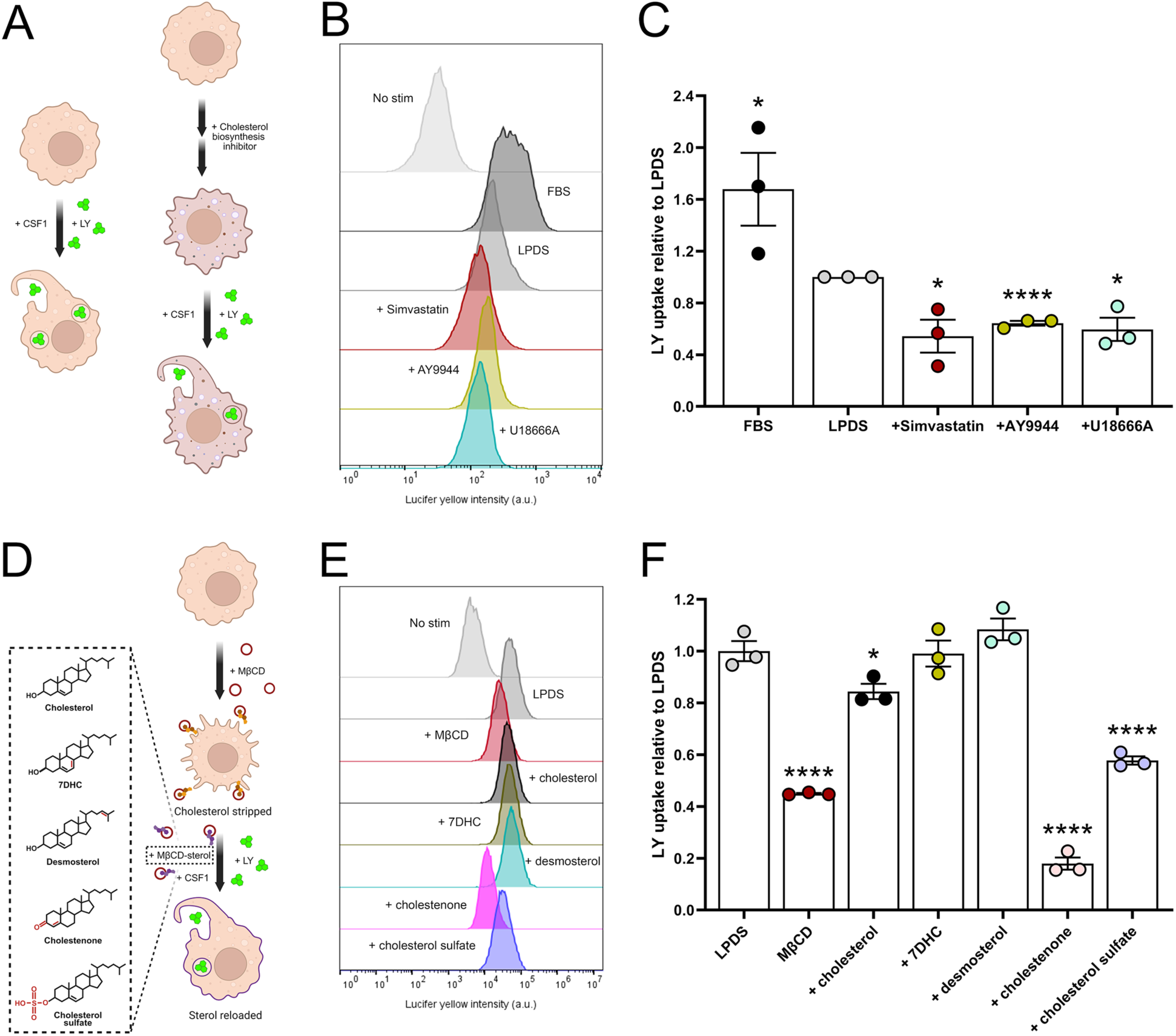
Macropinocytosis is regulated by both total sterol levels and sterol-mediated membrane phase separation. (A) Illustration of experimental design for analysis of cholesterol-biosynthesis inhibition impacts on macropinocytosis in BMDMs. (B) Representative histograms showing LY expression in BMDMs after cholesterol-targeted treatment paradigms. (C) Quantified LY uptake in BMDMs following impaired cholesterol biosynthesis (mean ± SEM; n = 3 biological replicates from 3 independent experiments). One-way ANOVA (F_4,10_ = 10.9, p < 0.0011) with Dunnett’s multiple comparisons test (*p ≤ 0.05 compared to LPDS culture). Unpaired t-test (LPDS vs +Simvastatin: t_4_ = 3.595, *p ≤ 0.05; LPDS vs +AY9944: t_4_ = 18.82, p < 0.0001; LPDS vs +U18666A: t_4_ = 4.525, *p ≤ 0.05). (D) Illustrated experimental design for acute modification of plasma membrane sterol content through cholesterol removal with MβCD followed by replacement with sterols of interest. Inset shows sterol structural differences (in red) relative to cholesterol. (E) Representative histograms of LY uptake in BMDMs after cholesterol removal and reloading of distinct sterols. (F) Quantified LY uptake following cholesterol removal and reloading of distinct sterols which support phase separation (cholesterol, 7DHC, or desmosterol) versus non-supportive sterols (cholestenone or cholesterol sulfate) (mean ± SEM; n = 3 biological replicates from 3 independent experiments). One-way ANOVA (F_6,14_ = 106.8, p < 0.0001) with Sidak’s multiple comparisons test (*p ≤ 0.05, ****p < 0.0001 compared to LPDS treatment).

We next questioned whether macrophage endocytic functions were dependent on sterol biochemistry impacts on phase separation within the plasma membrane (27). To do this, we acutely stripped cholesterol from the plasma membrane using methyl-β-cyclodextrin (MβCD) and replaced it with structurally distinct sterols which do or do not support phase separation (**Fig. 3D**). MβCD-treated BMDMs displayed a ∼50% reduction in LY uptake (**Fig. 3E, 3F**). Replacement of membranous cholesterol with cholesterol, 7DHC, or desmosterol rescued the reduction in LY uptake elicited by MβCD alone (**Fig. 3E, 3F**). However, neither cholestenone, where the C3 hydroxyl is replaced with carbonyl, nor cholesterol sulfate, in which the C3 hydroxyl is replaced with sulfate (**Fig. 3D**), were able to rescue LY uptake (**Fig. 3F**). These data suggest cholesterol content in macrophages is essential for proper endocytic processes, and that sterols which support membrane phase separation and organization are critical to macrophage function.

### Genetically compromised *Dhcr7* limits endocytic capabilities in macrophages

Since disruption of cholesterol homeostasis impeded critical endocytic processes in BMDMs (**Fig. 3**, supplemental Fig. S3), we analyzed functional activity in BMDMs derived from *Dhcr7*^T93M/+^ and *Dhcr7*^T93M/Δ3-5^ mice. While macropinocytosis was unaffected in BMDMs derived from *Dhcr7*^T93M/+^ and *Dhcr7*^T93M/Δ3-5^ mice maintained in cholesterol-rich FBS conditions (supplemental Fig. S4), LY uptake was inhibited in both *Dhcr7*^T93M/+^ and *Dhcr7*^T93M/Δ3-5^ BMDMs in LPDS conditions (**Fig. 4A, 4B**). To determine if macropinocytic function could be restored by biochemical rescue, BMDMs were returned to a cholesterol-rich environment (FBS conditions) or cholesterol was directly loaded into the plasma membrane (MβCD-cholesterol). While control macrophages (*Dhcr7*^+/+^) demonstrated a 30% increase in LY uptake after FBS supplementation and a 15% improvement after addition of MβCD-cholesterol (**Fig. 4C, 4D**), only direct loading of cholesterol with MβCD promoted macropinocytosis in *Dhcr7*^T93M/Δ3-5^ BMDMs (**Fig. 4C, 4D**). These findings further demonstrate the critical requirement for sterol homeostasis in macrophage function with disease relevance for SLOS.

**Figure 4.**
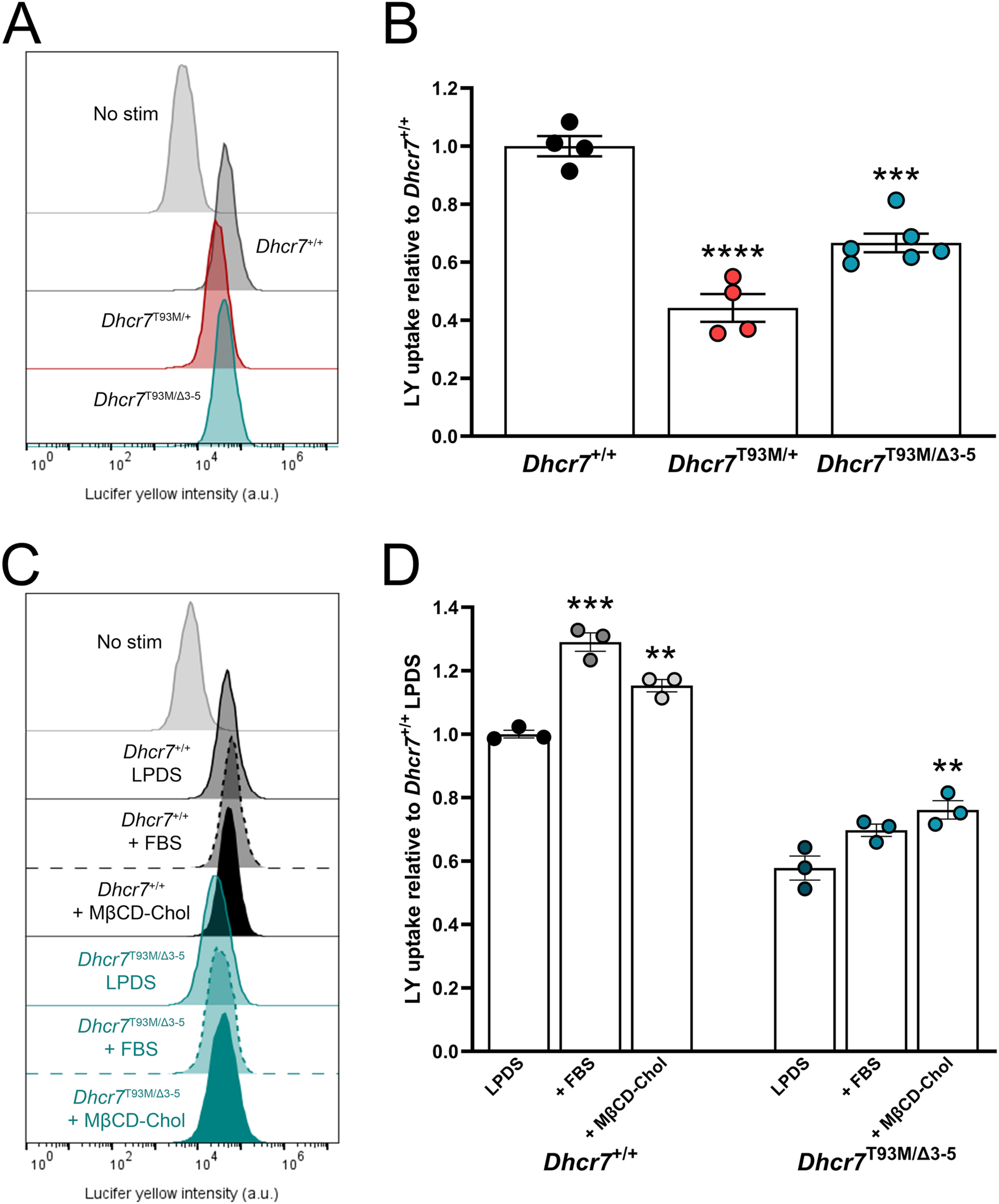
Genetic disruption of *Dhcr7* impairs macropinocytosis in macrophages. (A) Representative histograms of LY uptake in *Dhcr7^+/+^*, *Dhcr7*^T93M/+^, and *Dhcr7*^T93M/β^ BMDMs. (B) Quantified LY uptake in wild-type and *Dhcr7* mutant BMDMs (mean ± SEM; n = 4-6 biological replicates from 4 independent experiments). One-way ANOVA (F_2,11_ = 47.67, p < 0.0001) with Dunnett’s multiple comparisons test (***p ≤ 0.001; ****p < 0.0001 compared to *Dhcr7^+/+^*). (C) Representative histograms of BMDM LY uptake including replacement of LPDS with FBS or cholesterol-loaded MβCD (+MβCD-chol). (D) Quantified LY internalization in response to cholesterol-repletion in wild-type and *Dhcr7* mutant BMDMs (mean ± SEM; n = 3 biological replicates from 3 independent experiments). One-way ANOVA (*Dhcr7*^+/+^: F_2,6_ = 46.78, p ≤ 0.0002; *Dhcr7*^T93M/Δ3-5^: F_2,6_ = 9.777, p ≤ 0.0129) with Dunnett’s multiple comparisons test (**p ≤ 0.01, ***p ≤ 0.001 compared to *Dhcr7^+/+^* LPDS).

### Transcriptomic analyses reveal immunogenic and endosomal dysregulation in macrophages subsequent to impaired cholesterol biosynthesis

To uncover the mechanisms underlying the phenotypic and functional deficits observed in macrophages exhibiting sterol biochemistry restructuring, BMDMs challenged with cholesterol biosynthesis inhibition were analyzed by mRNA sequencing. The greatest number of differentially expressed genes was found in comparisons of BMDMs in FBS versus LPDS conditions (**Fig. 5A, 5B**), indicating macrophages rely on sterols sourced from their environment. Macrophages are specialized at detecting changes in environmental conditions (42), and BMDMs exposed to lipid-deficient (LPDS) relative to cholesterol-rich (FBS) media conditions exhibited dramatic transcriptome changes (2821 up, 2768 down). Differentially expressed genes from targeted cholesterol biosynthesis treatments (simvastatin, AY9944, and U18666A) versus LPDS conditions were mostly, but not exclusively, unique to the pharmacological agent utilized (**Fig. 5A, 5C**). Ingenuity pathway analyses of differentially expressed genes revealed both unique and similar processes impacted by the different treatment conditions. A major shared pathway altered across treatment was chemokine signaling (**Fig. 5D-F**, supplemental Fig. S5A, S5B), though AY9944 exclusively resulted in high expression of *Csf1* (**Fig. 5F**). These results point to a dysregulation in chemoregulatory responses associated with cholesterol homeostasis in BMDMs.

**Figure 5.**
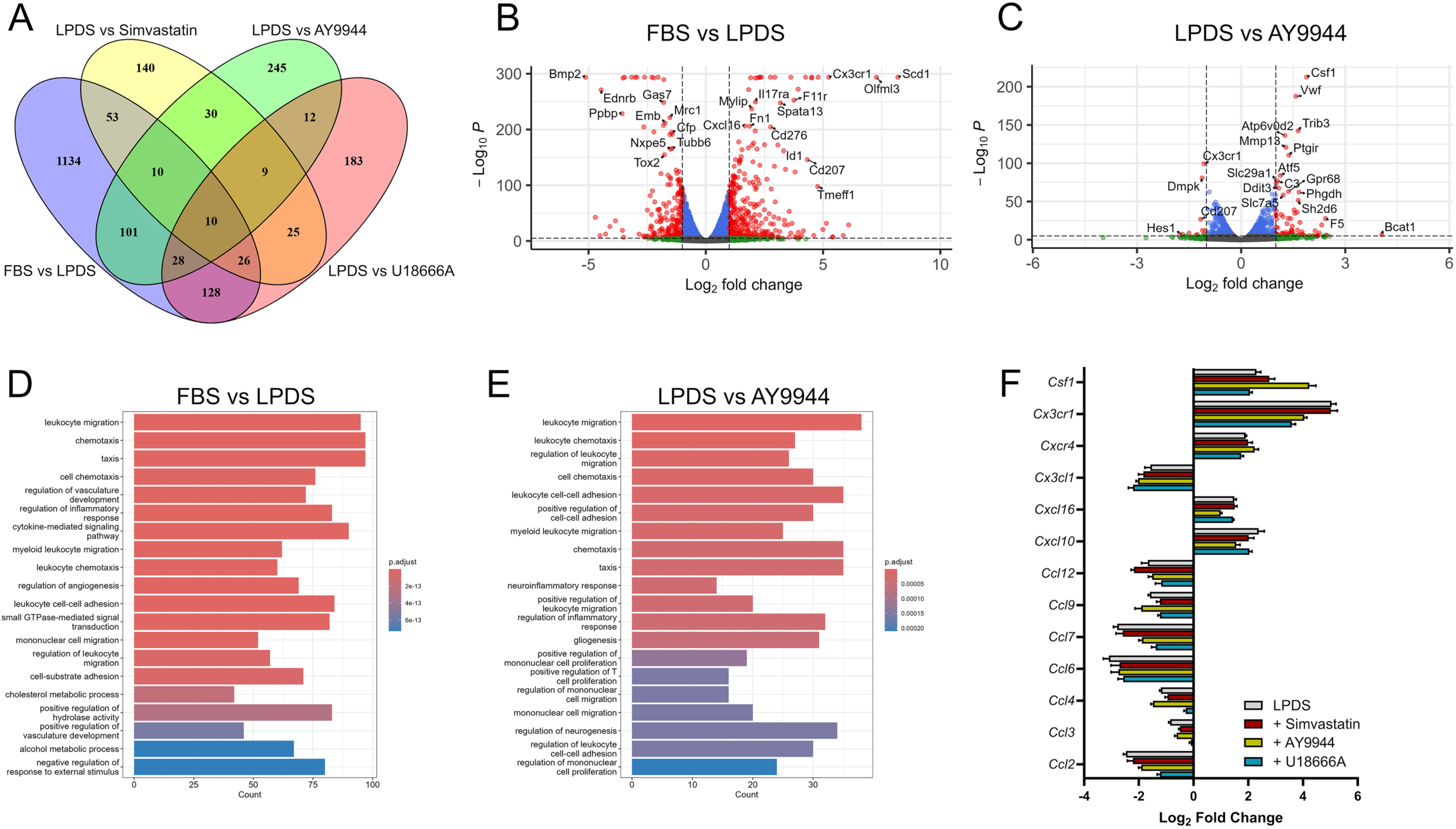
Sterol homeostatic change induces both shared and biochemistry-distinct cellular signaling deficits in macrophages. (A) Venn diagram illustrating differentially expressed transcripts that are shared and distinct between cholesterol-depleted conditions (n = 4 biological replicates per condition; displays genes for p-values < 0.05, log_2_ fold change > ±0.5). (B) Volcano plot indicating differentially expressed transcripts LPDS to FBS conditions (n = 4 biological replicates per condition). Red data points indicate differentially expressed transcripts (*p* < 0.05, log_2_ fold change ≥ ±0.5). (C) Volcano plot indicating differentially expressed transcripts when comparing LPDS to AY9944 treated conditions (n = 4 biological replicates per condition). Fold changes correspond to AY9944 expression relative to LPDS. Red data points indicate differentially expressed transcripts (*p* < 0.05, log_2_ fold change ≥±0.5. (D) Pathway analysis comparing LPDS to FBS conditions reveals alterations in immune response behavior in BMDMs (n = 4 biological replicates per condition). (E) Pathway analysis comparing AY9944 to LPDS conditions shows enhanced BMDM immunoreactive profile after AY9944 treatment (n = 4 biological replicates per conditions). (F) Transcriptional changes associated with chemokine signalling are significantly altered across treatments in BMDMs (mean ± SEM; n = 4 biological replicates per condition).

Further analyses of cholesterol-metabolic pathways revealed overall increased expression within genes of pre-squalene and post-squalene cholesterol biosynthesis, though gene expression was dampened by U18666A (supplemental Fig. S6A, S6B), suggesting possible desmosterol-mediated suppression of sterol synthesis. Relative to FBS, all treatment groups exhibited increased transcription of genes related to both low-density lipoprotein (LDL) receptor and lysosomal cholesterol transport (supplemental Fig. S6C). Genes encoding proteins related to cholesterol efflux, including the ATP-binding cassette transporter subfamilies A, B, and G exhibited reduced expression across treatments (supplemental Fig. S6D).

To examine if specific sterol biochemistries produced unique transcriptional networks within macrophages, we further analyzed common and unique transcriptional changes across treated BMDMs relative to LPDS (supplemental Fig. S7A, S7B). We first used GO analysis of transcripts that were common across treatment groups (78 genes total) to identify a shared immune signature (supplemental Fig. S7C). Activated immune-associated genes of interest included but *Il7r* (interleukin 7 receptor), *Il11ra1* (interleukin 11 receptor, alpha subunit), and *Ifrd1* (interferon related development regulator 1) (supplemental Fig. S7D). We next performed GO cellular component analysis on genes common between ≥2 treatment groups (supplemental Fig. S7E). Changes in gene expression shared between all three groups were related to early endosome signaling and cell surface expression (supplemental Fig. S7E). Comparisons between groups highlighted various pathways including ER signaling, endocytic/lysosomal signaling, and vesicular trafficking (supplemental Fig. S7E, S7F). These results demonstrate the transcriptomes of macrophages are dramatically altered by cholesterol biosynthesis inhibition and highlight sterol impacts on signaling networks linked to immune function and vesicular/endosomal transport.

### Impaired cholesterol production initiates digestive pathway in BMDMs

The transcriptomic profiles of macrophages with impaired cholesterol biosynthesis suggest there could be a dysfunction in immune responsivity in conjunction with vesicular trafficking defects. Interferon gamma (IFNγ) and lipopolysaccharide (LPS) polarized macrophages may take on a phagocytic phenotype from which endosomal and lysosomal processes become more engaged (43). Analysis of the early endosome maker Rab5 revealed BMDMs cultured in LPDS increased endosome labeling, an effect that was enhanced with TASIN-1 or AY9944 (supplemental Fig. S8A, S8B). Further analysis revealed increased expression of the late endosome/lysosome protein Rab7 after treatments with cholesterol biosynthesis inhibitors simvastatin, 17α-OHP, TASIN-1, or AY9944 (**Fig. 6A, 6B**). While inhibition of sterol esterification with avasimibe also increased Rab7 expression, the Dhcr24 antagonist U18666A had no effect on Rab7 expression (**Fig. 6B**). CD68 is a lysosomal protein that shuttles to the plasma membrane and binds to modified LDL-cholesterol and phosphatidylserine; CD68 expression increases in atherogenic conditions or LPS exposure (44). CD68 exhibited elevated expression with all treatments (**Fig. 6A, 6C**, supplemental Fig. S8, Fig. S9). Phalloidin staining also exhibited increased signal intensity after impairing cholesterol metabolism, suggesting excessive actin polymerization of defects in depolymerization (**Fig. 6A, 6D**, supplemental Fig. S8, Fig. S9).

**Figure 6.**
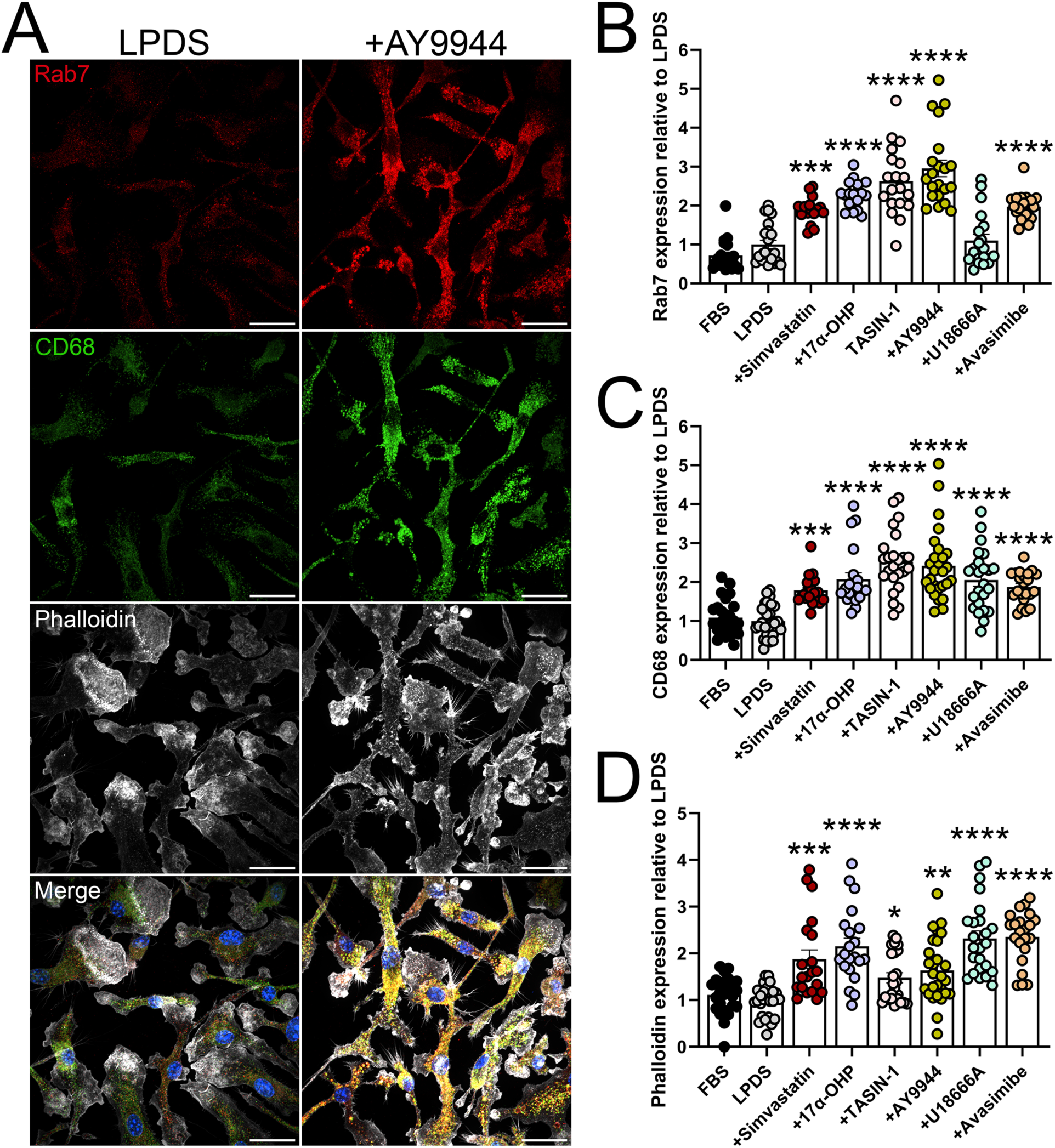
Sterol disruption induces immunoreactivity and morphological changes in macrophages. (A) Representative images for late endosomes/lysosomes (Rab7), immunoreactivity (CD68), and F-actin filaments (phalloidin-647) in BMDMs after 48 h exposure to LPDS or AY9944 treatment. Scale bar, 25 µm. (B) Simvastatin, 17α-OHP, TASIN-1, AY9944, and avasimibe promote accumulation of Rab7-positive late endosomes/endosomes (mean ± SEM; n = 16-23 images taken from 2 independent experiments). One-way ANOVA (F_7,146_ = 34.55, p < 0.0001) with Dunnett’s multiple comparisons test (***p ≤ 0.001; ****p < 0.0001). (C) Quantified CD68 expression in cholesterol metabolism inhibited BMDMs (mean ± SEM; n = 20-28 images taken from 3 independent experiments). One-way ANOVA (F_7,183_ = 19.38, p < 0.0001) with Dunnett’s multiple comparisons test (***p ≤ 0.001; ****p < 0.0001). (D) Quantified phalloidin-647 expression observed after disruption of cholesterol metabolism. One-way ANOVA (F_7,183_ = 16.71, p < 0.0001) with Dunnett’s multiple comparisons test (*p ≤ 0.05; **p ≤ 0.01; ****p < 0.0001).

The engulfment and storage of extracellular lipids is an important function of macrophages to prevent environmental stress and toxicity (45, 46). To determine if increased immunoreactivity due to cholesterol depletion impaired macrophage lipid storage capacity, we analyzed BMDMs for lipid droplet expression. Oleic acid (OA) supplementation increased neutral BODIPY 505/515 and perilipin-2 (PLN-2) staining in cells cultured in either FBS or LPDS conditions (supplemental Fig. S10). PLN-2 intensity was minimally increased with cholesterol synthesis inhibition (supplemental Fig. S11A, S11B), though the number of lipid droplets did not change relative to LPDS conditions (supplemental Fig. S11A, S11C). These data demonstrate that altered transcriptional program and cell surface protein expression upon cholesterol-deficient culture does not affect macrophage lipid storage mechanisms but is instead connected to dysfunctional lipid trafficking.

### Macrophages with genetic disruption of *Dhcr7* display reactive phenotype

To further define the relationship between immune activation and vesicular defects in response to cholesterol biosynthesis inhibition, we analyzed BMDMs from mice exhibiting mutations within *Dhcr7*, mimicking the cholesterol biosynthesis disorder SLOS (23, 24). While BMDMs from *Dhcr7* mutant mice (*Dhcr7*^T93M/+^, *Dhcr7*^T93M/Δ3-5^) exhibited lower levels of cholesterol relative to controls (*Dhcr7*^+/+^), there was no expected accumulation of the sterol precursor 7DHC (**Fig. 7A**) as previously observed in other cell types derived from these mice (22, 24, 47, 48). Since previous studies have demonstrated interferon signaling in response to pathogen exposure inhibits cholesterol biosynthesis (17, 18), we analyzed the impact of BMDM activation on sterol biosynthesis. Through stimulation of BMDMs with IFNγ, LPS (49), and AY9944 co-administration, IFNγ/LPS stimulation eliminated the accumulation of sterol precursors in AY9944-treated BMDMs and prevented cholesterol loss below LPDS conditions (**Fig. 7B**). These results demonstrate that sterol biochemical changes in *Dhcr7* macrophages are dependent upon immune responsivity, preventing buildup of cholesterol intermediates.

**Figure 7.**
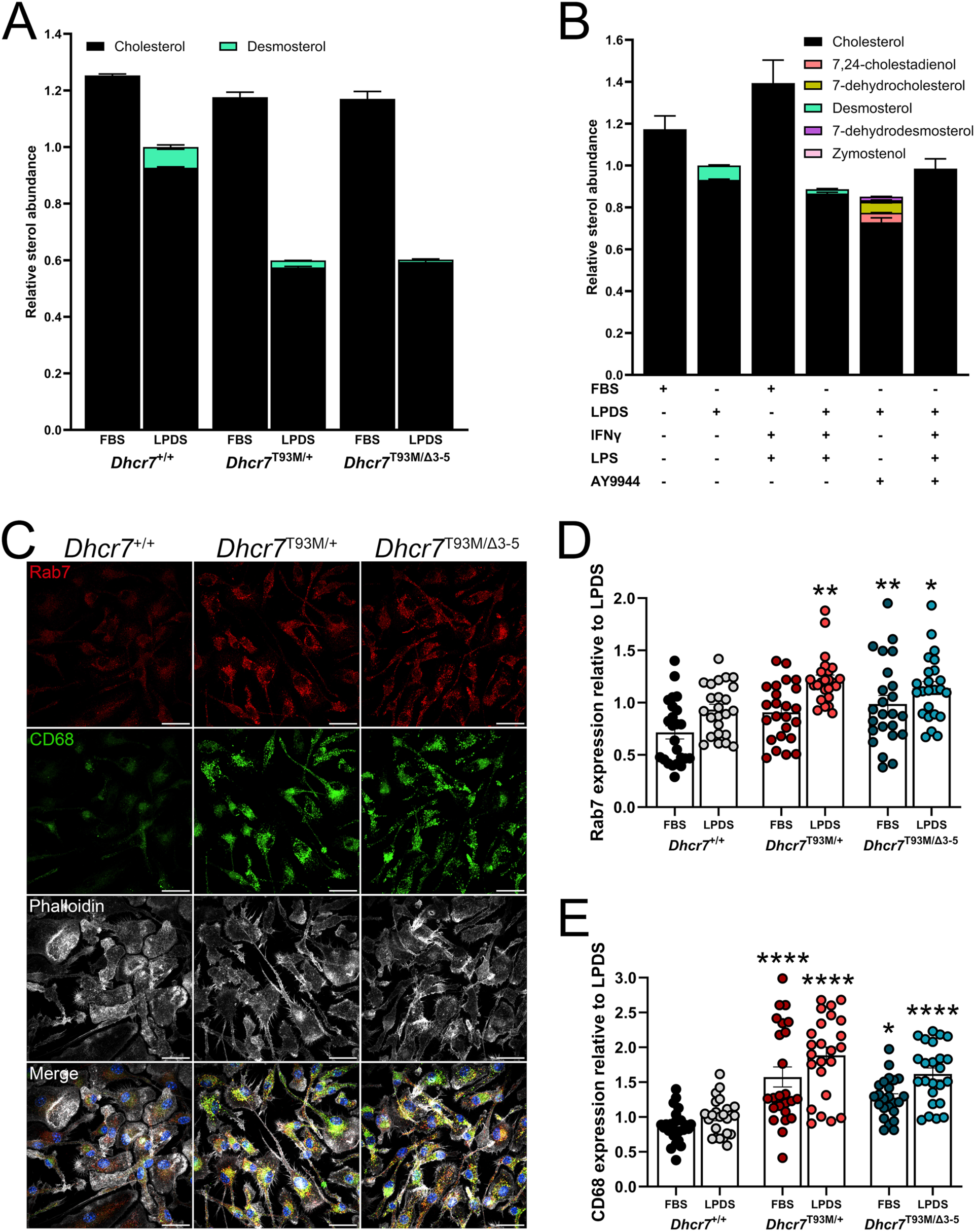
Biochemical profiles in macrophages from *Dhcr7* deficient mice are driven by increased immunoreactivity. (A) Quantified GC-MS analyses of BMDMs from *Dhcr7*^T93M/+^ and *Dhcr7*^T93M/β3-5^ show reduced cholesterol but no 7DHC accumulation (mean ± SEM; n = 2 biological replicates from 2 independent experiments). (B) Quantified GC-MS analyses of BMDMs treated with AY9944 alongside IFNγ/LPS (mean ± SEM; n = 2 biological replicates from 2 independent experiments). (C) Representative images of Rab7 and CD68 from *Dhcr7*^+/+^, *Dhcr7*^T93M/+^, and *Dhcr7*^T93M/β^ BMDMs in LPDS conditions. Scale bars, 25 µm. (D) Quantified Rab7 expression in wild-type and *Dhcr7* mutant BMDMs (mean ± SEM; n = 24 images taken from 3 independent experiments). Two-way ANOVA (media effect: F_1,138_ = 20.65, p < 0.0001; genotype effect: F_2,138_ = 10.02, p < 0.0001) with Sidak’s multiple comparisons test (*p ≤ 0.05, **p ≤ 0.01 relative to *Dhcr7^+/+^* LPDS). (E) Quantified CD68 expression in *Dhcr7^+/+^* versus *Dhcr7* mutant (*Dhcr7*^T93M/+^, *Dhcr7*^T93M/β^) BMDMs (mean ± SEM; n = 24 images taken from 3 independent experiments). Two-way ANOVA (media effect: F_1,138_ = 12.34, p ≤ 0.0006; genotype effect: F_2,138_ = 38.94, p < 0.0001) with Sidak’s multiple comparisons test (*p ≤ 0.05, ****p < 0.0001 relative to *Dhcr7^+/+^* LPDS).

We next assessed if *Dhcr7* mutant BMDMs exhibit accumulation of endosomal/lysosomal markers, as in AY9944-treated pharmacological models. Both *Dhcr7*^T93M/+^ and *Dhcr7*^T93M/Δ3-5^ BMDMs expressed higher levels of Rab7 relative to control BMDMs (**Fig. 7C, 7D**, supplemental Fig. S12). Additionally, we found increased CD68 expression in *Dhcr7*^T93M/+^ and *Dhcr7*^T93M/Δ3-5^ BMDMs in both FBS and LPDS conditions (**Figure 7C, 7E**, supplemental Fig. S12). These findings indicate that genetic disruption of cholesterol biosynthesis causes immunological and vesicular defects in macrophages.

## DISCUSSION

Our work demonstrates a critical connection between sterol metabolism and macrophage cell function and signaling. From an evolutionary perspective, cholesterol likely better supports the complex array of signaling pathways observed in mammalian systems compared to intermediate sterols due to improved membrane structural integrity and lipid ordering (50). Our study addresses the functional importance of precursor sterol molecules on macrophages and membrane biology to extend previous findings. For example, desmosterol is a natural ligand for the liver x receptor which mediates expression of cholesterol efflux and cholesterol synthesis genes in macrophages; however, desmosterol may play additional roles in other cell types (51, 52). Outside of cholesterol production, 7DHC serves as a precursor to vitamin D, and the degradation of DHCR7 in the presence of high cholesterol and desmosterol can enhance vitamin D production (53).

Macrophages rely on changes in membrane fluidity to engage phagocytic behavior (54). Both phagocytosis and macropinocytosis require significant membrane reformations through recruitment of lipids and cytoskeletal elements to distinct cellular loci (55–57). In our study, the substitution of cholesterol with desmosterol or 7DHC in the plasma membrane rescued impaired macropinocytosis (**Fig. 3C, 3D**). We previously showed reconstitution of membrane sterols with desmosterol or 7DHC also restored clathrin-mediated endocytosis (27). Lipid rafts are an important feature for membrane bending processes, like those related to endocytosis (58, 59). However, desmosterol’s double bond at C24 is thought to impair lipid ordering, preventing it from replacing cholesterol in lipid rafts (60). Our macropinocytosis rescue experiment suggests membrane sterols that retain the 3β-hydroxyl group support macropinocytosis (**Fig. 3C, 3D**). While sterol loading may artificially impact lipid distribution between the endoplasmic reticulum and the plasma membrane, affecting cholesterol sensing and transport mechanisms (61), further studies into the impact of defined sterols on endocytic processes are necessary to define this concept.

Though cholesterol depletion impaired macrophage endocytosis through macropinocytosis and clathrin-mediated endocytosis (**Fig. 3**, supplemental Fig. S3), we saw higher expression of endosomal and lysosomal markers after sterol biosynthesis inhibition (**Fig. 6**, supplemental Fig. S8, Fig. S9). Vesicular sorting and digestive pathways were also dysfunctional in macrophages from a model of SLOS (**Fig. 7**, supplemental Fig. S8) and were differentially expressed in our transcriptomic analyses (supplemental Fig. S7E, S7F). Phagocytic cells undergo lysosomal restructuring during endocytic product digestion, which could be stimulus-induced (62). Disruption of normal sorting pathways may be attributed to several factors, including changes to vesicle acidification (63, 64), autophagy-induced redirection of endocytic cargo (65), or impaired vesicle fusion (66). The inability of cholesterol to be transported out of lysosomes also disrupts macrophage intracellular vesicular transport processes (67). We have recently shown that cholesterol starvation interferes with endosomal sorting pathways, causing accumulation of late endosomes which fuse with autophagosomes (29). It is possible this is represented within the results here, where reduced cholesterol, and not the presence of sterol intermediates, initiates vesicular trafficking irregularities. Importantly, regulation of lysosomal function is directly linked to immune responsivity in macrophages (68) and chemotactic response (69), which we observed to be heavily impacted with cholesterol biosynthesis disruption (**Fig. 2**, **Fig. 5D-F**). Additional studies are necessary to more thoroughly detail these connections.

Macrophage cholesterol metabolism was previously demonstrated to be dependent on cellular immunogenicity (14). Cellular redistribution of cholesterol through metabolic reorganization regulates IFN signaling (18). Previous work also reported that impaired regulation of sterol biosynthetic pathways was initiated by type I IFN response (17). Following cholesterol biosynthesis inhibition, we found differential expression of various immunomodulatory signaling mechanisms (supplemental Fig. S7C, S7D). Within our transcriptomic data, we found altered expression of type I IFN signaling genes such as *Ifrd1* shared between all cholesterol synthesis treatments (supplemental Fig. S7D). Combined with the absence of intermediate sterol accumulation in *Dhcr7*^T93M/+^ and *Dhcr7*^T93M/Δ3-5^ macrophages (**Fig. 7A, 7B**), our work is in line with previous studies linking macrophage immune activation with sterol synthesis and metabolism (17, 18). Our data also suggest macrophages from *Dhcr7*^T93M/+^ and *Dhcr7*^T93M/Δ3-5^ are likely immunoactivated prior to isolation in response to unknown stimuli. Since cholesterol-rich lipid rafts house pattern recognition receptors to initiate immune responses to pathogens (70) and perturbations to lipid ordering at these sites can impair receptor signaling (71), our findings suggest sterol-specific regulation of intracellular lipid organization may contribute to macrophage biology. Previous studies in SLOS models demonstrated innate immune system pathways are disrupted (72, 73). Additional work to define the *in vivo* drivers of immune activation in *Dhcr7* models also require further investigation.

The relationship between genetic disorders of cholesterol biosynthesis and immune system dysregulation remains underexplored. Within models of SLOS, we previously demonstrated heightened immunoreactivity in *Dhcr7* deficient astrocytes seems to be driven by macrophage-like microglia (22). More recent work also showed that both *Dhcr7*^T93M/Δ3-5^-derived macrophages and pharmacological inhibition of *Dhcr7* in macrophages induced immune activation (72, 73), including altered *Tnf* and *Cxcl* signaling as we show here (**Fig. 5F**). Clinically, cholesterol supplementation to SLOS subjects was suggested to limit infections (74), though the mechanism of action and direct impacts on immune cell function and signaling are unknown. However, the extent to which disrupted immune function is evident across different tissues and genetically distinct disorders of cholesterol biosynthesis is unknown. Cholesterol biosynthesis disorders can impact multiple organ systems, including nervous, immune, epidermal, and skeletal (3). Whether disrupted macrophage function contributes to skeletal abnormalities within SLOS, Conradi-Hünermann, or other cholesterol biosynthesis disorders, for example, requires further study (7, 75). While we demonstrate a connection between immune-associated anomalies and compromised cholesterol biosynthesis, additional studies are necessary to fully elucidate this relationship in the context of human disease.

## CONCLUSION

In summary, we have delineated a role for disrupted cholesterol homeostasis and sterol biosynthesis within immune cell function and signaling. Investigation of endocytic processes showed severe deficiencies with cholesterol depletion, but replenishing membrane sterols with distinct structural features restored function. Further, macrophages from SLOS mouse models displayed atypical sterol profiles, which were likely due to IFN-promoted sterol reorganization and impaired macropinocytosis. Concomitantly, these results further elucidate the critical role of lipid homeostasis for immune cell function and suggest immune cell dysfunction may contribute to the pathology of cholesterol biosynthesis disorders.

## DATA AVAILABILITY

All data are provided within the main text and supplemental document. Raw RNA sequencing data are freely available at RNA sequencing data are available in the Gene Expression Omnibus (GEO) database (http://www.ncbi.nlm.nih.gov/gds) under the accession number GSE300253. Further information and requests for resources and reagents should be directed to and will be fulfilled, if possible, by the lead contact, Dr. Kevin Francis.

## ACKNOWLEDGEMENTS

Thank you to Forbes Porter (*Eunice Kennedy Shriver* National Institute of Child Health and Human Development) for sharing of *Dhcr7* mouse models. Thank you to Adam Hoppe (South Dakota State University) for helpful discussions. We would like to thank the Center for Brain and Behavior Research at the University of South Dakota for supporting this project. We would like to thank the following core facilities at Sanford Research for experimental support: Imaging, Flow Cytometry, and Functional Genomics and Bioinformatics cores. Illustrations were created using BioRender (https://biorender.com/).

Any opinions, findings and conclusions expressed in this material are those of the author(s) and do not necessarily reflect the views of the National Institutes of Health.

## Abbreviations

lipoprotein-deficient serum, LPDS; bone marrow-derived macrophage, BMDM; Smith-Lemli-Opitz syndrome, SLOS; 7-dehydrocholesterol, 7DHC; quantitative real-time polymerase chain reaction, qRT-PCR; 17α-hydroxyprogesterone, 17α-OHP; Gene Ontology, GO; colony-stimulating factor 1, CSF1; paraformaldehyde, PFA; oleic acid, OA; methyl-β-cyclodextrin, MβCD; Lucifer yellow, LY

