## Supplemental Information for "Macrophage signaling and function are regulated by distinct sterol biochemistries"

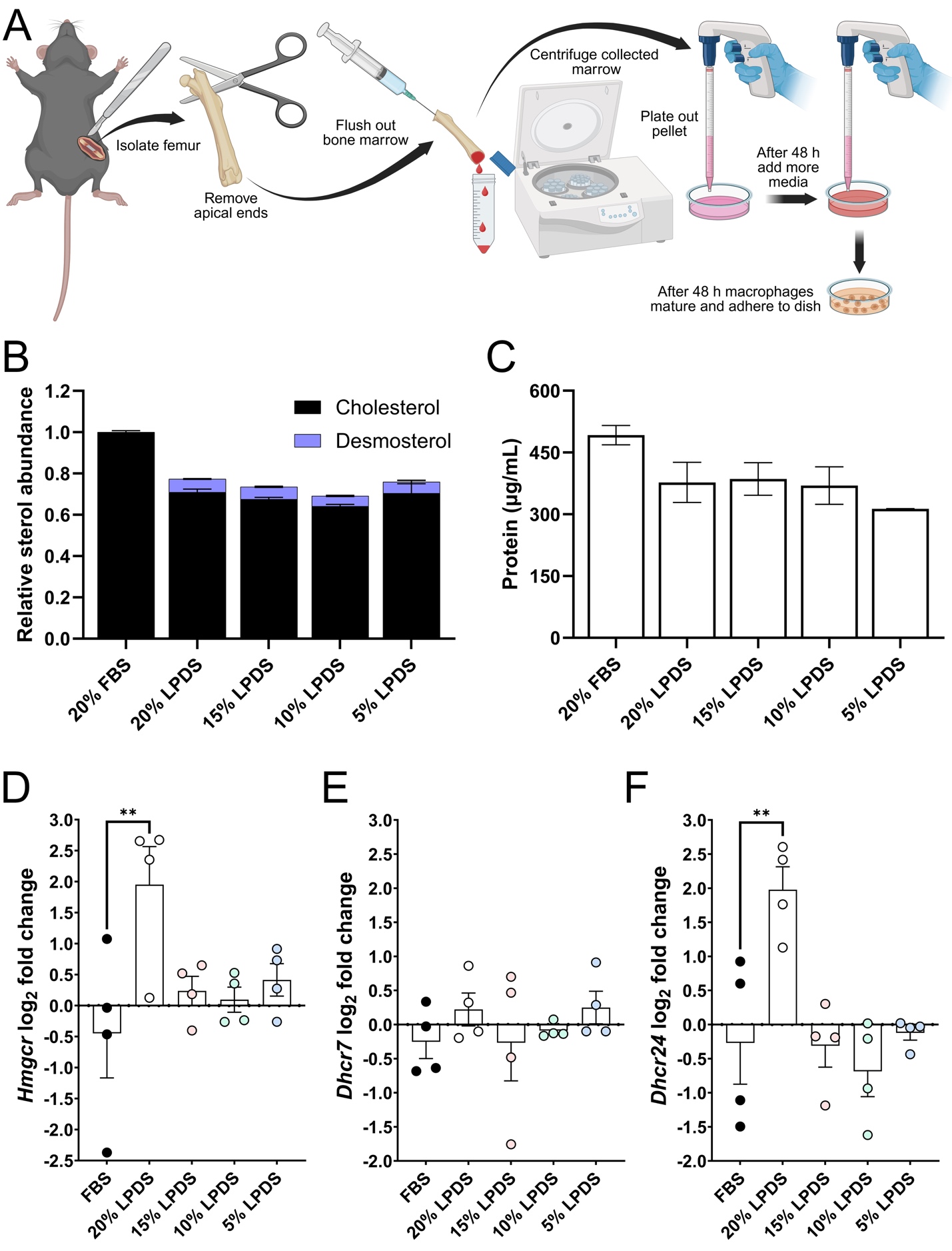

**Figure S1. Bone marrow-derived macrophages exhibit differential sterol profiles upon culture in lipoprotein deficient conditions. Related to Figure 1.**

(A) Illustration summarizing the protocol for BMDM isolation from mouse femur.
(B) BMDMs cultured in LPDS for 48 h at varying concentrations exhibit reduced cellular cholesterol and desmosterol accumulation (mean ± SEM; n = 2 biological replicates from 2 independent experiments).
(C) Normalized protein content in BMDMs after incubation in various LPDS concentrations for 48 h (mean ± SEM; n = 2 biological replicates from 2 independent experiments).
(D) Quantified *Hmgcr* expression in BMDMs in FBS versus LPDS conditions for 48 h (mean ± SEM; n = 4 biological replicates from 2 independent experiments). One-way ANOVA (F_4,15_ = 3.83, p ≤ 0.0244) with Dunnett’s multiple comparisons test (**p < 0.01 compared to FBS).
(E) Quantified *Dhcr7* levels in FBS versus LPDS conditions for 48 h (mean ± SEM; n = 4 biological replicates from 2 independent experiments). One-way ANOVA (F_4,15_ = 0.6399, p = 0.6421).
(F) Quantified *Dhcr24* expression in FBS versus LPDS conditions for 48 h (mean ± SEM; n = 4 biological replicates from 2 independent experiments). One-way ANOVA (F_4,15_ = 7.751, p ≤ 0.0014) with Dunnett’s multiple comparisons test (**p < 0.01 compared to FBS).

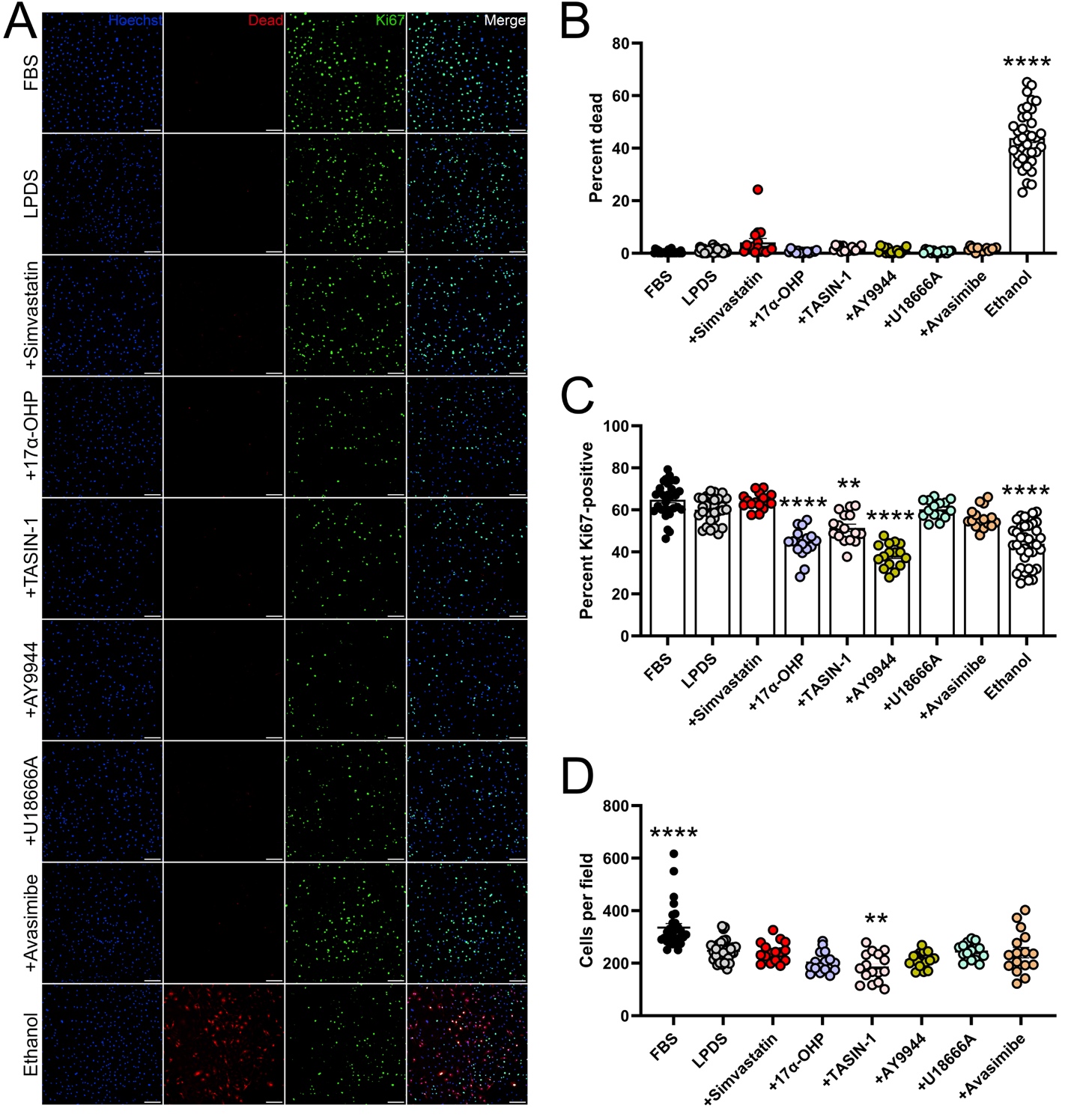
**Figure S2. Impaired cholesterol biosynthesis inhibits macrophage proliferation. Related to Figure 1.**

(A) Representative images of cell viability (red) and proliferative capacity (Ki67; green) in BMDMs. Hoechst counterstain is blue. Scale bars, 100 µm.
(B) Quantified cell death following inhibition of cholesterol biosynthesis for 48 h (mean ± SEM; n = 15-37 images taken from 3 independent experiments). One-way ANOVA (F_8,185_ = 270.1, p < 0.0001) with Dunnett’s multiple comparisons test (****p < 0.0001 compared to LPDS).
(C) Quantified percent Ki67 positive cells following inhibition of cholesterol biosynthesis for 48 h (mean ± SEM; n = 15-37 images taken from 3 independent experiments). One-way ANOVA (F_8,185_ = 40.21, p < 0.0001) with Dunnett’s multiple comparisons test (**p < 0.01; ****p < 0.0001 compared to LPDS).
(D) Quantified BMDMs per field when cultured with LPDS, FBS, or cholesterol biosynthesis inhibitors (mean ± SEM; n = 15-32 images taken from 3 independent experiments). One-way ANOVA (F_7,149_ = 16.21, p < 0.0001) with Dunnett’s multiple comparisons test (**p < 0.01; ****p < 0.0001 compared to LPDS).

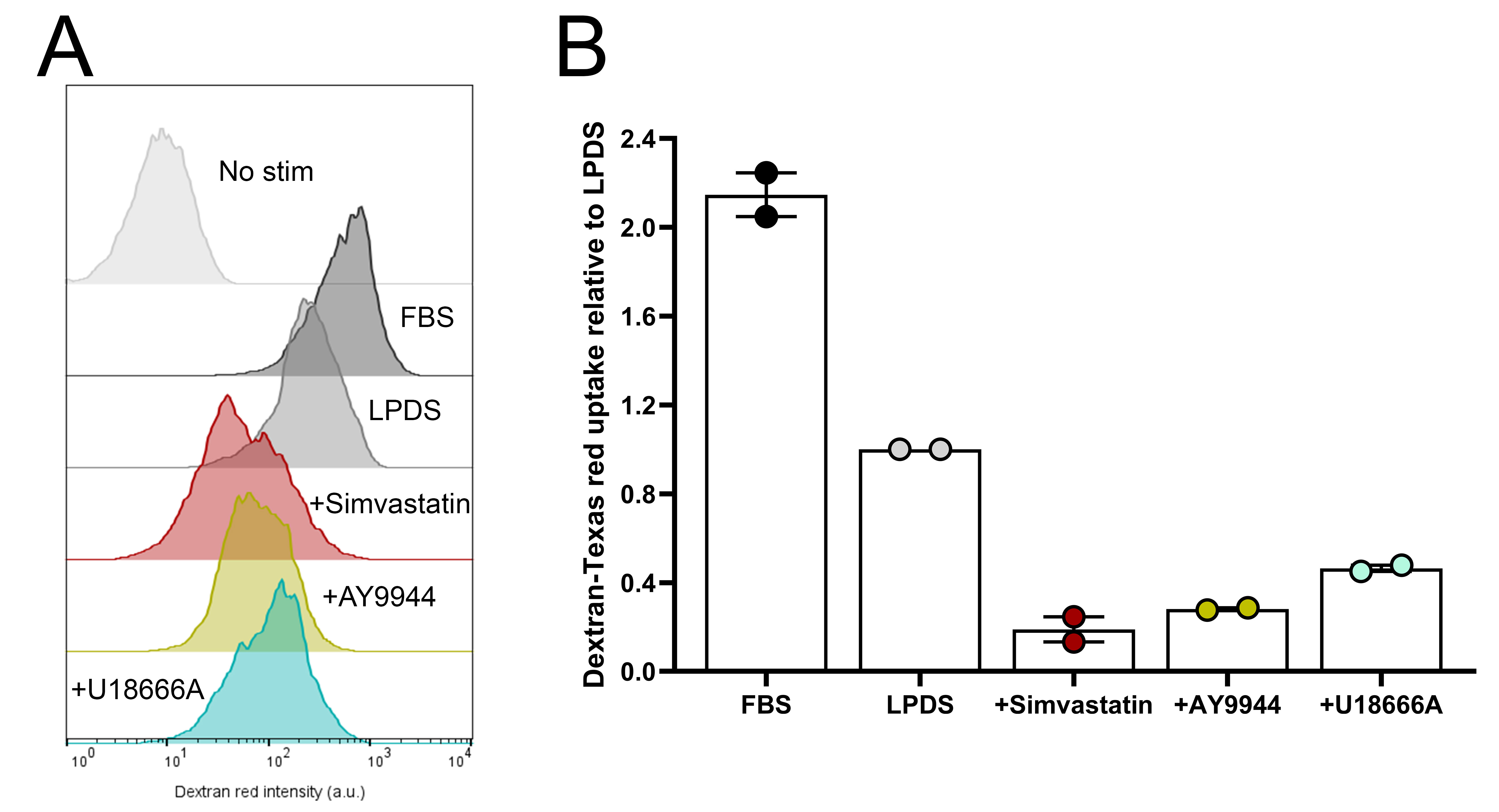

**Figure S3. Clathrin-mediated endocytosis is attenuated in macrophages upon cholesterol homeostasis loss. Related to Figure 3.**

(A) Representative histograms for Texas red dextran uptake in BMDMs in the presence or absence of cholesterol biosynthesis inhibition.
(B) Quantified Texas red dextran internalization after inhibition of cholesterol biosynthesis (mean ± SEM; n = 2 biological replicates from 2 independent experiments).

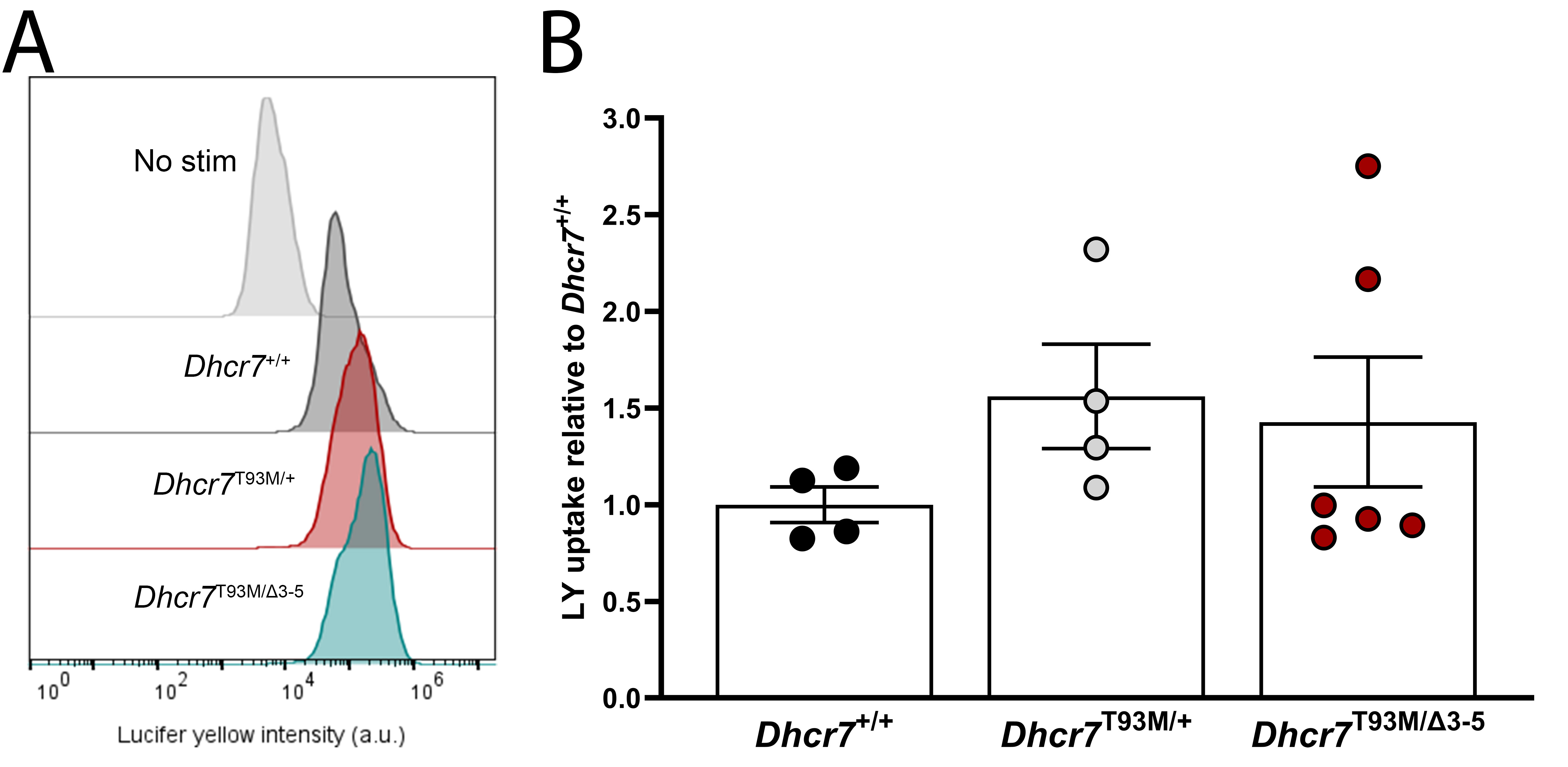

**Figure S4. Macrophages from mouse models of Smith-Lemli-Opitz syndrome exhibit normal macropinocytic function in cholesterol rich conditions. Related to Figure 4.**

(A) Representative histograms for LY uptake in *Dhcr7*^+/+^, *Dhcr7*^T93M/+^, and *Dhcr7*^T93M/Δ3-5^ BMDMs cultured in FBS conditions.
(B) Quantified LY uptake in *Dhcr7*^+/+^, *Dhcr7*^T93M/+^, and *Dhcr7*^T93M/Δ3-5^ BMDMs maintained in FBS conditions (mean ± SEM; n = 4-6 biological replicates from 4 independent experiment). One-way ANOVA (F_2,11_ = 0.8875, p = 0.4392).

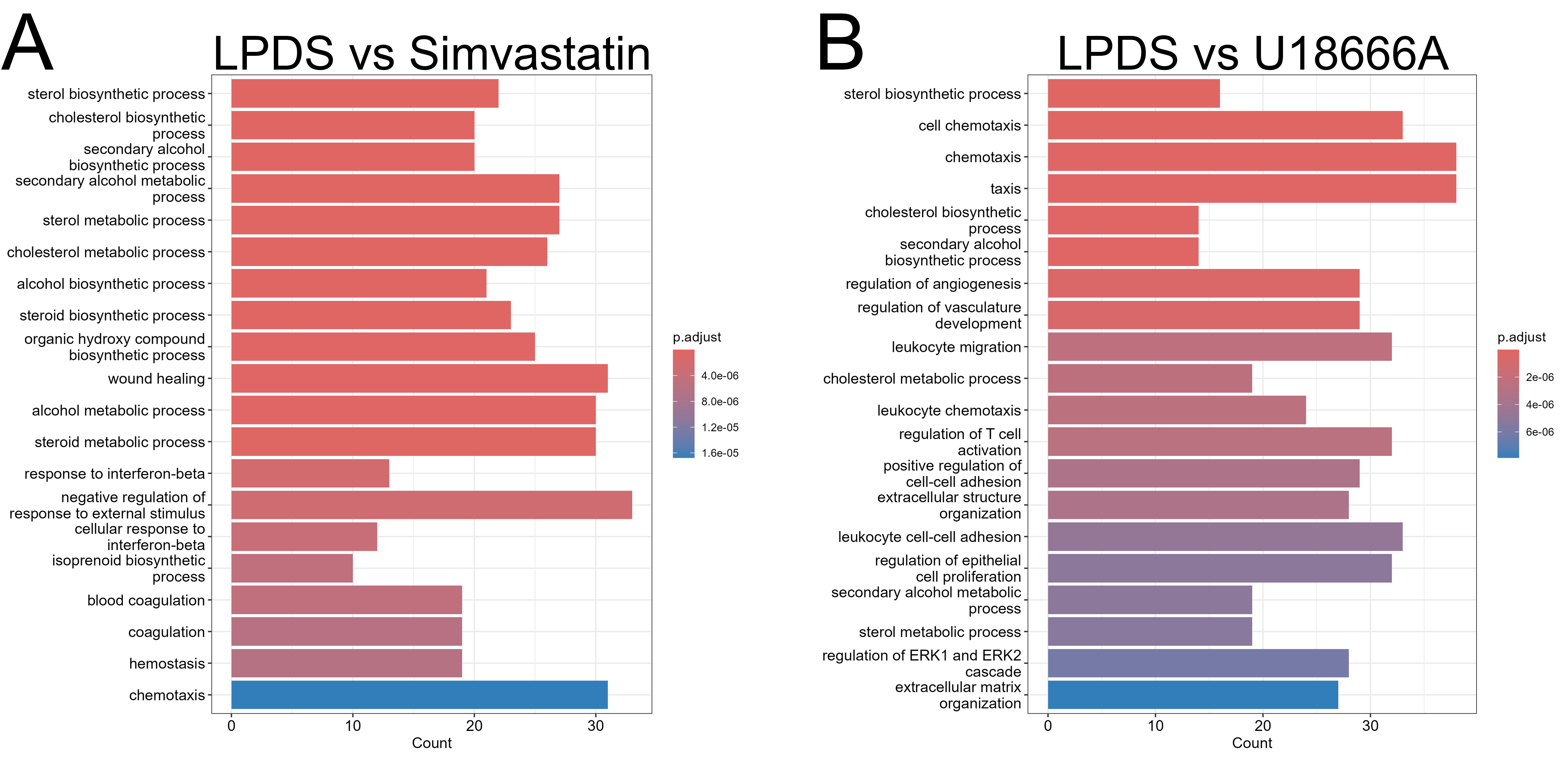

**Figure S5. Pathway analyses of cholesterol biosynthesis impacts on macrophage signaling. Related to Figure 5.**

(A) Pathway analysis of simvastatin treatment compared to LPDS conditions indicate altered sterol biosynthetic and immune response pathways (n = 4 biological replicates).
(B) Pathway analysis of U18666A treatment compared to LPDS culture revealed enhanced sterol biosynthetic and immune response after U18666A treatment (n = 4 biological replicates).

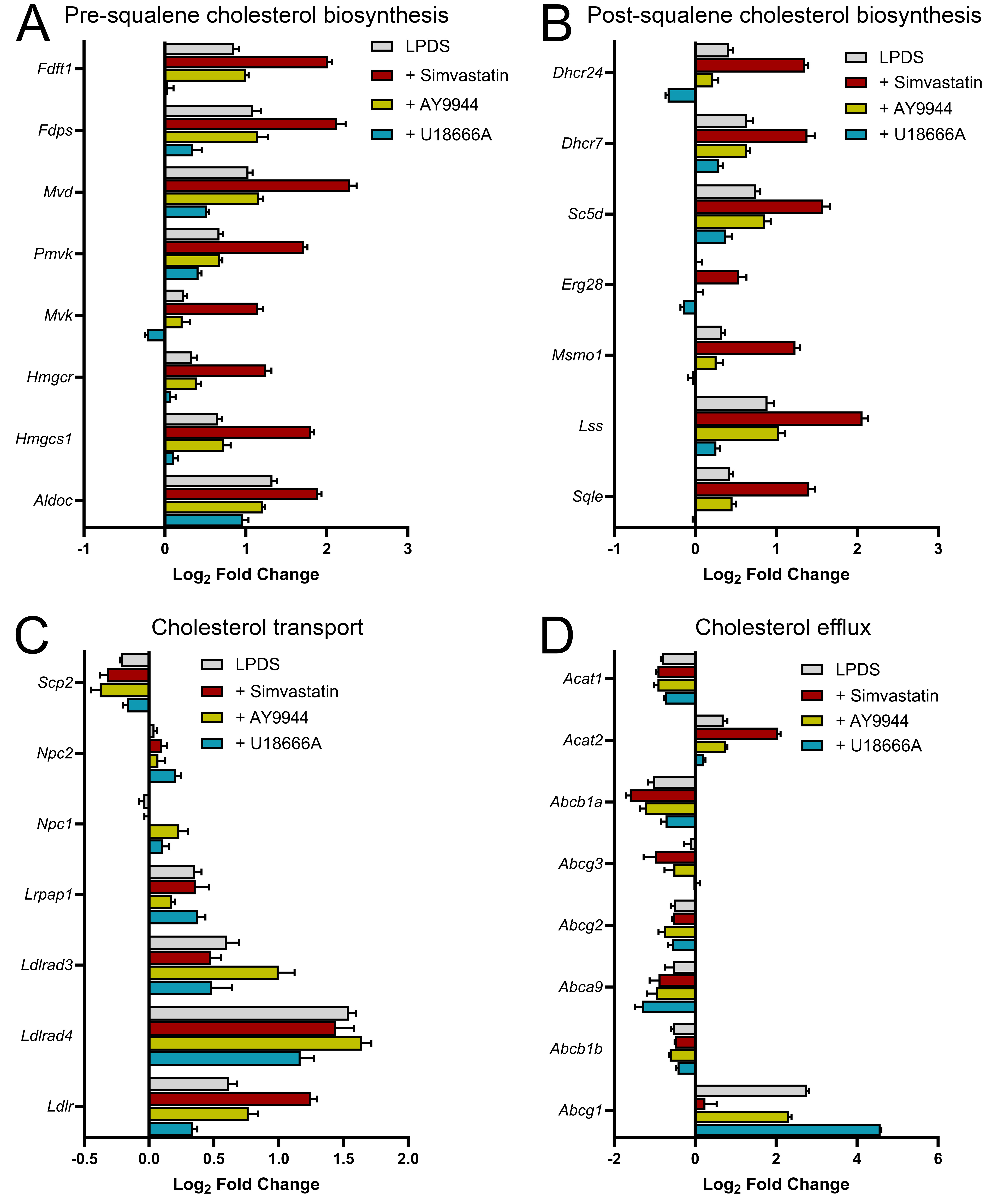
**Figure S6. Impaired cholesterol biosynthesis in macrophages alters expression of sterol metabolism-associated transcripts. Related to Figure 5.**

(A) Transcripts associated with activation of pre-squalene cholesterol biosynthesis are increased in treatments that reduce cholesterol biosynthesis (mean ± SEM; n = 4 biological replicates).
(B) Transcripts associated with post-squalene cholesterol biosynthesis are increased in conditions of impaired sterol metabolism (mean ± SEM; n = 4 biological replicates).
(C) Transcripts associated with regulation of cholesterol synthesis and transport are altered in response to cholesterol changes in macrophages (mean ± SEM; n = 4 biological replicates).
(D) Cholesterol deficient conditions induce transcriptional changes related to cholesterol efflux and lipid storage (mean ± SEM; n = 4 biological replicates).

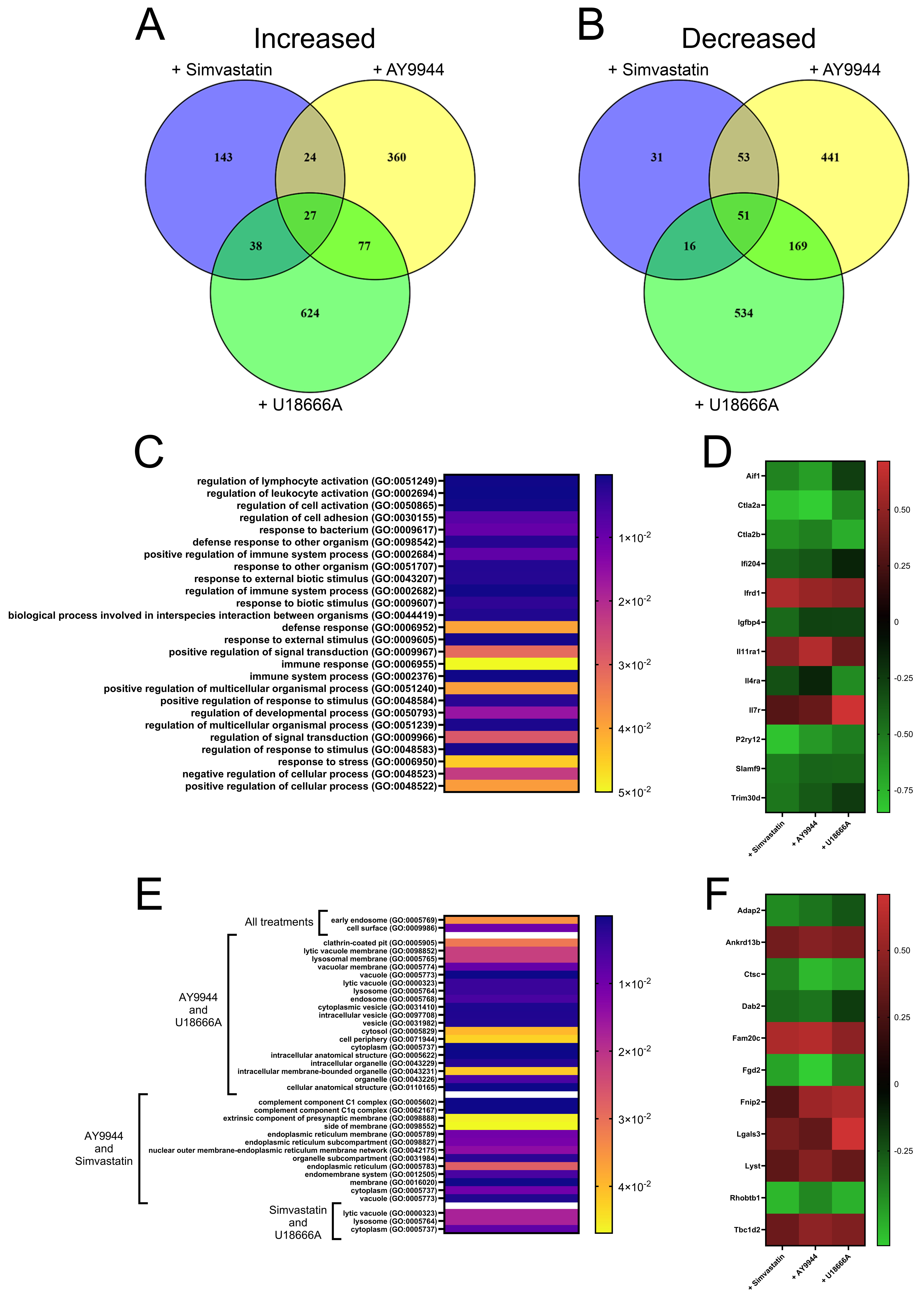

**Figure S7. Gene Ontology pathway analyses reveal impacts of cholesterol biosynthesis inhibition on macrophage immune responsivity and vesicular trafficking processes. Related to Figure 5.**

(A) Venn diagram illustrating transcripts significantly increased after cholesterol biosynthesis inhibition relative to LPDS conditions (n = 4 biological replicates per condition; log_2_ fold change ≥ ±0.5, *p* < 0.05).
(B) Venn diagram showing transcripts significantly decreased after disrupted cholesterol biosynthesis relative to LPDS conditions (n = 4 biological replicates per condition; log_2_ fold change ≥ ±0.5, *p* < 0.05).
(C) GO pathway analyses for biological processes of shared transcripts suggest immune-related dysfunction in macrophages treated with cholesterol biosynthesis inhibitors relative to LPDS conditions. (n = 4 biological replicates per condition).
(D) Heat map showing changes in selected transcripts related to immune response that are differentially expressed in simvastatin, AY9944, and U18666A treatments relative to LPDS control conditions (mean ± SEM; n = 4 biological replicates per condition).
(E) GO pathway analyses for cellular components of shared transcripts show disruption of vesicular trafficking pathways in macrophages treated with cholesterol biosynthesis inhibitors relative to LPDS conditions. (n = 4 biological replicates per condition).
(F) Heat map showing selected transcripts associated with intracellular vesicular trafficking that are differentially expressed in simvastatin, AY9944, and U18666A treatments relative to LPDS conditions (mean ± SEM; n = 4 biological replicates per condition).

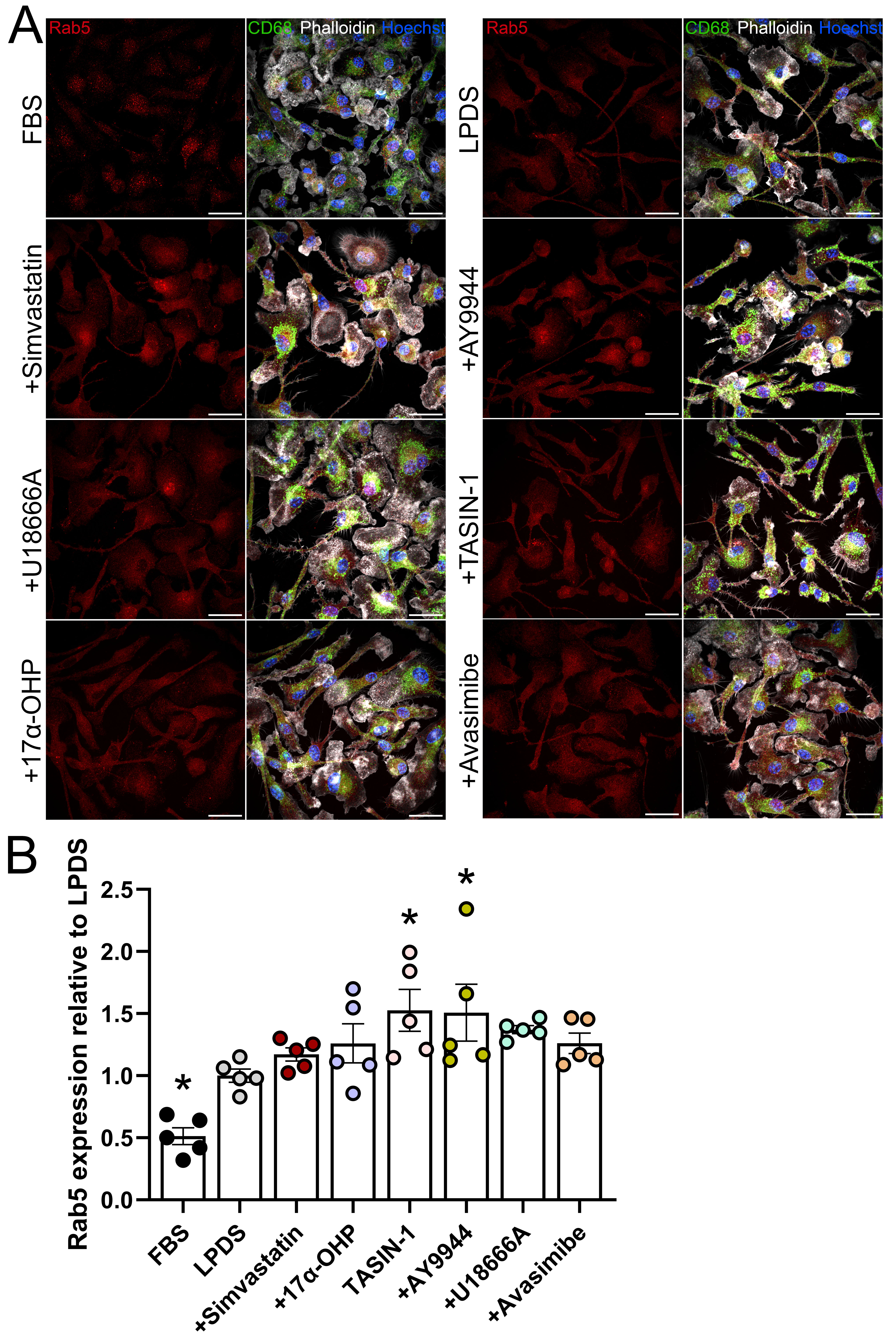

**Figure S8. Dysregulation of cholesterol biosynthesis impacts early endosomal pathways. Related to Figure 6.**

(A) Representative images of Rab5 expression with CD68, phalloidin-647, and Hoechst counterstain in BMDMs. Scale bars, 25 µm.
(B) Analysis of Rab5 expression after cholesterol synthesis antagonism relative to control conditions (mean ± SEM; n = 5 images taken from 1 independent experiment). One-way ANOVA (F_7,32_ = 6.921, p < 0.0001) with Dunnett’s multiple comparisons test (*p < 0.05 compared to LPDS conditions).

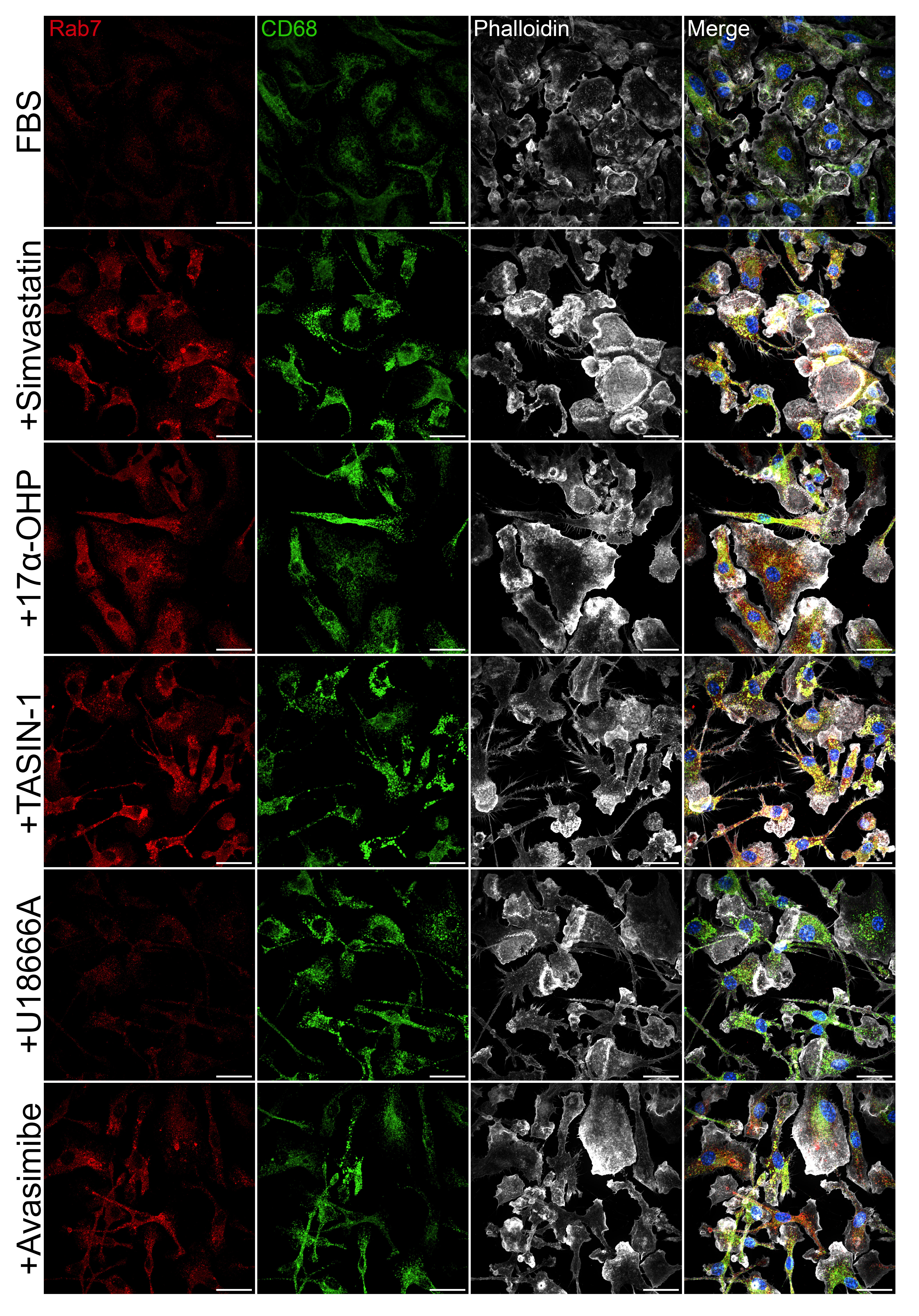
**Figure S9. Disruption of cholesterol metabolism promotes immunoreactivity in macrophages. Related to Figure 6.**

Representative images of markers of late endosomes/lysosomes (Rab7), immunoreactivity (CD68), and F-actin filaments (phalloidin) in BMDMs with or without cholesterol biosynthesis inhibition. Hoechst counterstain is blue. Scale bars, 25 µm.

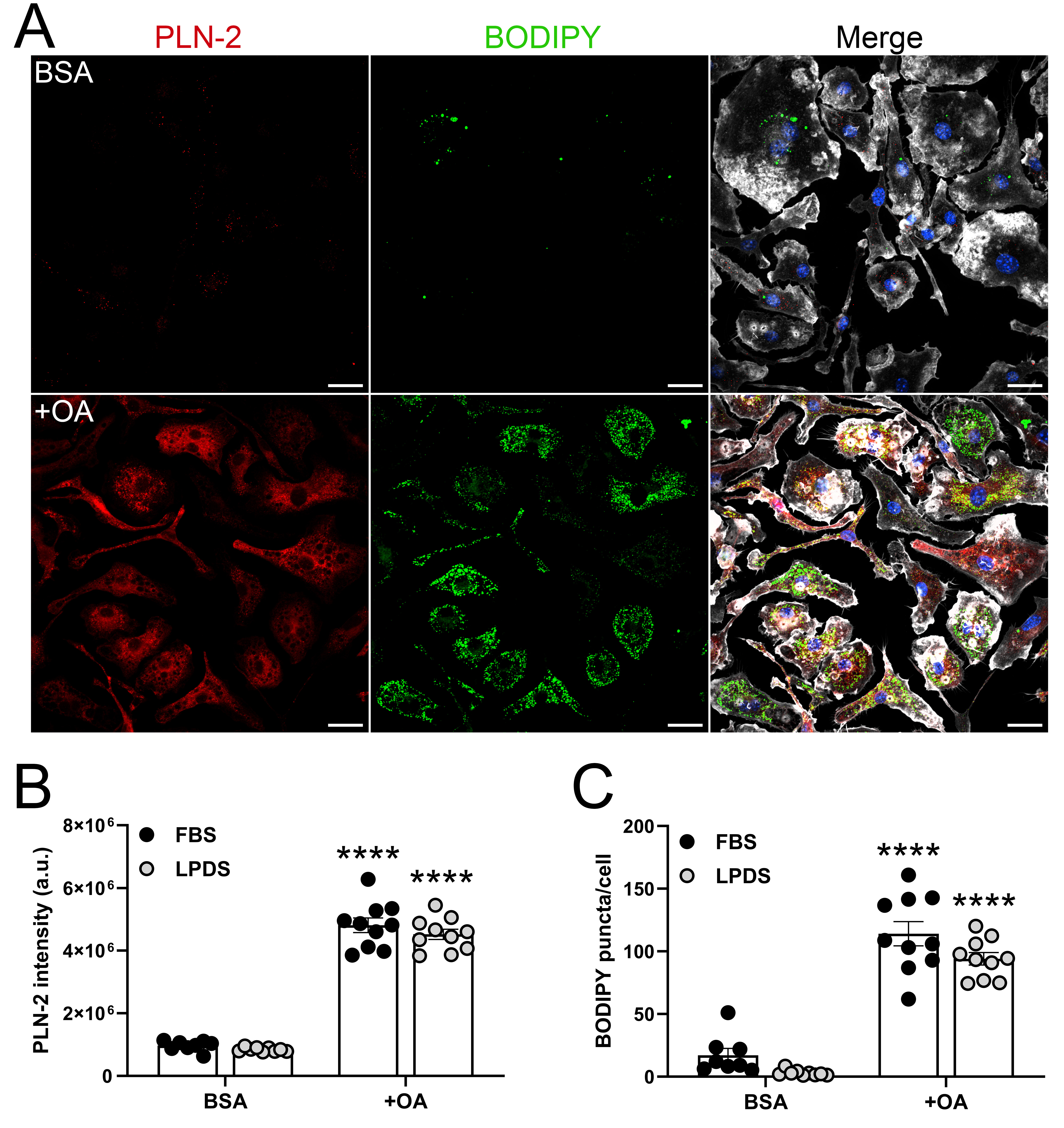
**Figure S10. Lipoprotein deficient culture does not prevent macrophage lipid storage. Related to Figure 6.**

(A) Representative images of PLN-2 and BODIPY 505/515 in LPDS cultured BMDMs after BSA or oleic acid (OA) treatment. Hoechst counterstain is blue. Scale bars, 25 µm.
(B) Quantified PLN-2 expression normalized by cell count after OA supplementation compared to BSA in FBS and LPDS conditions (mean ± SEM; n = 8-10 images taken from 2 independent experiments). Two-way ANOVA (Media effect: F_1,33_ = 571.6, p < 0.0001) with Tukey’s multiple comparisons test (****p < 0.0001 compared to BSA).
(C) Neutral lipid puncta labeled with BODIPY 505/515 are increased after OA treatment compared to BSA control conditions in both FBS and LPDS conditions (mean ± SEM; n = 8-10 images taken from 2 independent experiments). Two-way ANOVA (Media effect: F_1,33_ = 217.2, p < 0.0001; OA effect: F_1,33_ = 7.212, p < 0.0112) with Tukey’s multiple comparisons test (****p < 0.0001 compared to BSA).

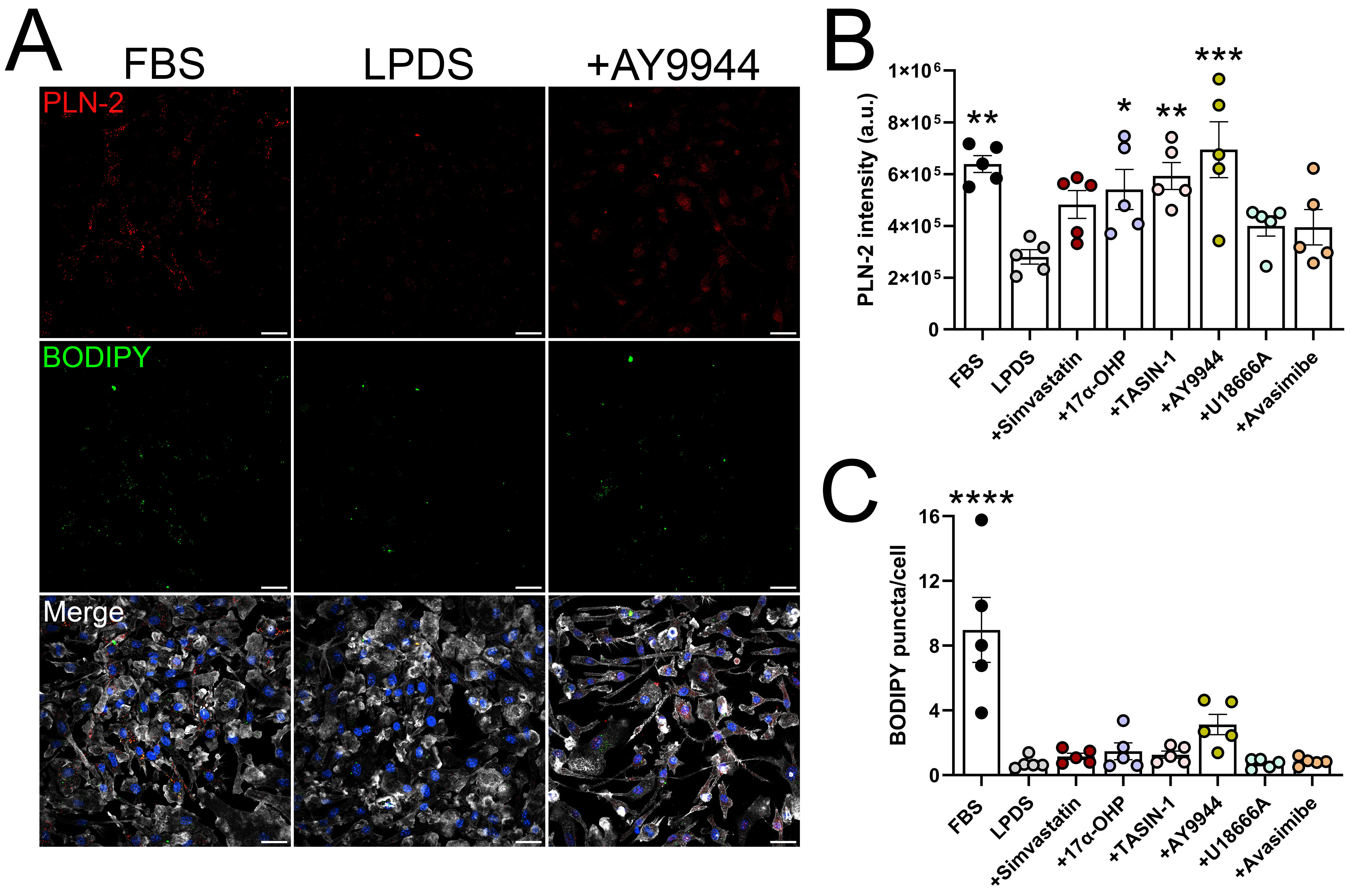

**Figure S11. Disrupted cholesterol biosynthesis does not promote lipid droplet formation in macrophages. Related to Figure 6.**

(A) Representative images of PLN-2 and BODIPY 505/515 in FBS, LPDS, or AY9944 cultured BMDMs. Scale bars, 25 µm.
(B) Quantified PLN-2 expression normalized by cell count in BMDMs with or without inhibition of cholesterol metabolism (mean ± SEM; n = 5 images taken from 1 independent experiment). One-way ANOVA (F_7,32_ = 5.006, p ≤ 0.0007) with Dunnett’s multiple comparisons test (*p ≤ 0.05; **p ≤ 0.01; ***p ≤ 0.001 compared to LPDS).
(C) Quantified BODIPY 505/515 puncta in BMDMs with or without inhibition of cholesterol metabolism (mean ± SEM; n = 5 images taken from 1 independent experiment). One-way ANOVA (F_7,32_ = 13.11, p < 0.0001) with Dunnett’s multiple comparisons test (****p < 0.001 compared to LPDS).

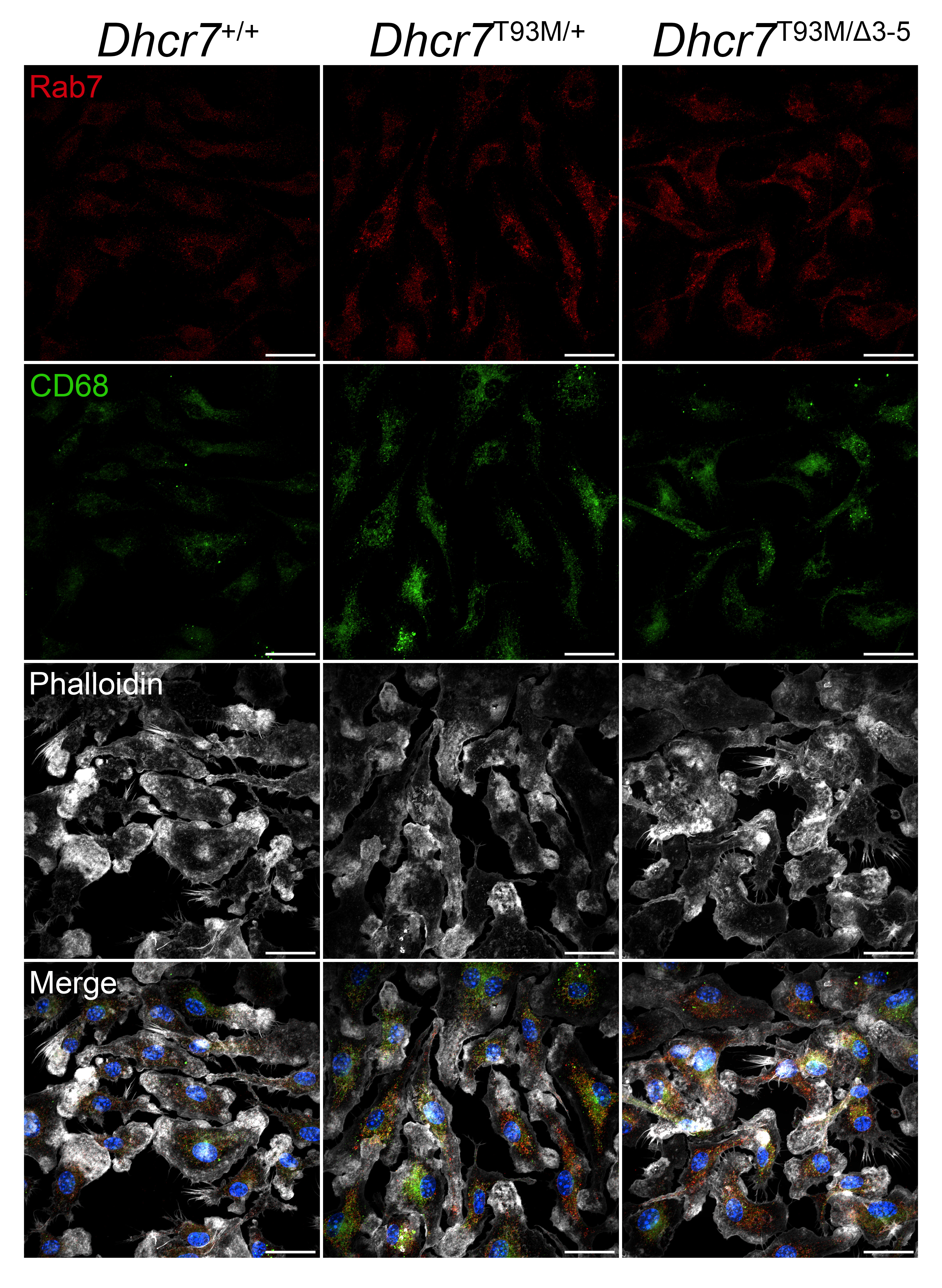

**Figure S12. Macrophages derived from mouse models of Smith-Lemli-Opitz syndrome display increased expression of markers of immunoreactivity. Related to Figure 7.**

Representative images of markers of late endosomes/lysosomes (Rab7) and immunoreactivity (CD68) in FBS-cultured BMDMs derived from wild-type (*Dhcr7*^+/+^) and mutant (*Dhcr7*^T93M/+^, *Dhcr7*^T93M/Δ3-5^) mice. Scale bar, 25 µm.
